# Ecological context unmasks cryptic effects of glyphosate tolerance on soybean metabolism and performance of the virus vector *Epilachna varivestis*

**DOI:** 10.64898/2026.03.18.712557

**Authors:** Hannier Pulido, Consuelo M. De Moraes, Mark C. Mescher

## Abstract

- Glyphosate-tolerant Roundup Ready (RR) soybean, engineered with the CP4 EPSPS gene, is one of the world’s most widely cultivated GM crops, yet its ecological consequences under realistic multi-species biotic conditions remain poorly understood. We investigated how RR soybean and its near-isogenic non-GM counterpart respond to concurrent colonization by two rhizobacteria, *Bradyrhizobium japonicum* and *Delftia acidovorans*, infection by Bean pod mottle virus (BPMV), and feeding by the virus vector *Epilachna varivestis*.
- Using a fully factorial multi-species design, we combined LC-MS/GC-MS metabolomics, weighted co-expression network analysis (WGCNA), and herbivore performance bioassays to assess genotype-dependent shifts in soybean metabolism and their consequences for herbivore performance.
- Under baseline conditions, RR and non-GM plants were metabolically indistinguishable. Under concurrent microbial and viral stress, RR plants diverged markedly, prioritizing selective isoflavonoid accumulation and lipid remodeling over broad-spectrum defenses, and showing attenuated rhizobacteria-mediated benefits for herbivore survival. These genotype-specific effects were entirely absent in single-species treatments.
- Transgene effects on soybean metabolism and tritrophic interactions are cryptic under simplified experimental conditions but emerge clearly under ecologically realistic multi-species stress, with direct implications for how GM crops are evaluated in agricultural and regulatory contexts.

## Introduction

Plant–microbe interactions can significantly influence plant metabolic pathways, thereby shaping ecological interactions with other organisms. Beneficial rhizobacteria, for instance, are known to regulate the synthesis of primary and secondary metabolites, as well as the release of volatile organic compounds, which can impact plant interactions with herbivores, their natural enemies, and vector-borne pathogens (Pineda *et al*., 2013; Pulido *et al*., 2019, 2024). Conversely, it is well established that pathogens can also alter plant phenotypes in ways that may facilitate their transmission by arthropod vectors (Eigenbrode *et al*., 2018). Since both mutualistic and pathogenic microorganisms can influence overlapping molecular and biochemical pathways—such as those governing plant defense and nutritional quality for herbivores—their concurrent presence in a plant host raises important ecological questions. In particular, their potentially conflicting effects on plant phenotypes could generate complex outcomes when plants are simultaneously colonized by both beneficial and harmful symbionts. Despite the growing body of research in this field, few studies have explored these interactions in the context of genetically modified (GM) crop varieties. Consequently, almost nothing is known about how GM crops integrate and respond to these kinds of concurrent multi-species interactions. Here, we investigate whether a glyphosate-tolerant soybean variety diverges from its near-isogenic non-GM counterpart in its responses to concurrent rhizobacterial colonization, viral infection, and specialist herbivory, and whether any such divergence reshapes tritrophic outcomes.

Soybean (*Glycine max*) is one of the world’s most widely cultivated crops, and glyphosate-tolerant Roundup Ready (RR) varieties — engineered with the CP4 EPSPS gene from *Agrobacterium tumefaciens* — account for the majority of global production. In soybean, the native EPSPS gene encodes an enzyme that functions in the penultimate step of the shikimic acid pathway, catalyzing a reaction between shikimate-3-phosphate and phosphoenolpyruvate to produce enolpyruvylshikimate-3-phosphate (Garg *et al*., 2014). This compound is used to produce aromatic amino acids and other aromatic compounds, including flavonoids that serve many important roles as pigments and signaling molecules mediating plant-microbe interactions. Glyphosate functions by inhibiting the native EPSPS gene, but transgenic plants expressing a modified EPSPS gene maintain EPSPS function under glyphosate applications (Imran *et al*., 2017). Because the EPSPS enzyme sits at a critical branchpoint feeding carbon into aromatic amino acids, flavonoids, and defense-related phenylpropanoids, modifications to this enzyme have the potential to alter how plants prioritize metabolic investment under stress, including stress imposed by interacting microbes and herbivores. Importantly, such effects may not be apparent under baseline conditions but could emerge when the shikimate pathway is actively recruited during biotic challenge.

Studies utilizing advanced analytical platforms, such as gas chromatography-mass spectrometry (GC-MS) and liquid chromatography-mass spectrometry (LC-MS), have generally reported minimal differences in metabolite profiles between GM and conventional soybeans, with variations often falling within the natural variability observed across cultivars and environmental conditions (Clarke *et al*., 2013; Kusano *et al*., 2015). For instance, McCann et al. (2005) identified no statistically significant alterations in primary metabolites (e.g., amino acids, organic acids) or secondary metabolites (e.g., isoflavones) attributable to the CP4 EPSPS transgene, reinforcing the principle of substantial equivalence. Yet the RR cultivar has been shown to affect biological nitrogen fixation (Hungria *et al*., 2014) and soil microbial community structure (Liu *et al*., 2005), suggesting that its ecological consequences are not fixed properties of the genotype but emerge from interactions with the biotic environment. This pattern motivates explicitly multi-species experimental designs.

These studies have focused primarily on how GM modifications alter plant-associated microbial communities but have rarely examined the downstream consequences of these alterations for higher-level species interactions (Silva *et al*., 2021). In agricultural ecosystems, soybean plants interact simultaneously with beneficial and pathogenic microorganisms that collectively modulate plant metabolism and stress responses (Riviezzi *et al*., 2021; Pulido *et al*., 2024) — and how GM modifications shape plant responses within such multi-species contexts remains largely unexplored.

While extensive studies have examined the effects of glyphosate application on RR soybean-rhizobia symbiosis (Zablotowicz & Reddy, 2007), less attention has been given to how the RR transgene itself might influence these interactions in the absence of herbicide. This gap is notable given the crucial roles these microorganisms play in soybean productivity and health (Giesler *et al*., 2002; Fonseca de Souza *et al*., 2025). The EPSPS enzyme is pivotal for production of aromatic compounds involved in primary metabolism, defense, and flavonoid signaling, all of which also mediate plant interactions with bacteria and viruses. We predict that RR plants, which have a different version of the EPSPS gene, will exhibit unique metabolic responses to viral and bacterial symbionts that propagate upward to modify interactions with a key specialist herbivore. To explore this prediction, we studied changes in herbivore suitability and metabolites in near isoline RR and nonRR genotypes during interactions with *Bradyrhizobium japonicum*, a key nitrogen-fixing symbiont, and bean pod mottle virus (BPMV) (*Comovirus siliquae*, family *Secoviridae*), a significant soybean pathogen. Building on prior work (Pulido *et al*., 2024) we also studied how inoculation with a non-specialist plant growth-promoting rhizobacteria (*Delftia acidovorans*) influences RR and nonRR soybean herbivore quality and metabolites, both after single inoculation and in combination with rhizobia and virus. By embedding the GM comparison within a fully factorial multi-species design, spanning mutualistic rhizobacteria, a viral pathogen, and a specialist herbivore, our study tests whether the ecological consequences of the RR transgene are context-dependent, and whether they emerge only when plants are challenged by the kinds of concurrent biotic interactions they face in agricultural fields.

We exposed RR soybean and its near-isogenic NonRR counterpart to single and combined colonization by *Bradyrhizobium japonicum* (Bj) and *Delftia acidovorans* (Da), with and without BPMV infection, and assessed resulting changes in leaf metabolomes (LC-MS/GC-MS, WGCNA) and in the performance of larvae of the specialist beetle herbivore *Epilachna varivestis*. We predicted that, under baseline conditions, RR and NonRR plants would be metabolically similar, but that GM-specific effects would emerge under multi-species biotic stress reflecting altered integration of signals from co-occurring mutualists and pathogens through a modified shikimate pathway. We further predicted that any such genotype-specific metabolic shifts would carry upward to influence herbivore performance in ways not predictable from single-species treatments alone.

## Methods

### Rhizobacteria and BPMV culture conditions

*B. japonicum* and *D. acidovorans* cultures were obtained from a commercial inoculant (BrettYoung) and maintained long-term as glycerol stocks (30%) at -80°C under sterile conditions. To inoculate bacteria, sterile techniques were used to revive stocks in yeast-mannitol broth incubated at 25°C under shaking conditions—40 hours for *D. acidovorans* and 80 hours for *B. japonicum*. Prior to inoculation, bacterial suspensions were adjusted to a final concentration of 1×10⁹ CFU/ml. Three-day-old soybean seedlings were inoculated by pipetting 1 ml of the prepared bacterial suspension directly into the soil. Control plants received an equivalent volume of sterile broth medium without bacteria. The BPMV strain used in this study was originally collected from soybean fields in Ohio (USA), purified to isolate virions, and provided by Dr. Peg Redinbaugh.

### Plant growth and experimental design

We implemented a multi-factorial experimental design to examine the individual and combined effects of *Delftia acidovorans* (Da), *Bradyrhizobium japonicum* (Bj), and BPMV on Roundup Ready (RR) soybean and its near-isogenic non-GM counterpart (NonRR), allowing us to assess plant responses under single, dual, and triple colonization conditions.

Soybean seeds from the MON89788 glyphosate-tolerant cultivar and its near-isogenic non-GM counterpart were provided by Monsanto. Seeds were surface-sterilized by immersion in a 10% sodium hypochlorite solution for 5 minutes, rinsed thoroughly with ultrapure water, and germinated in autoclaved, mycorrhiza-free growing medium (Premier Pro-Mix, Griffin Supplies) at 120°C for 40 minutes. After three days, seedlings were transplanted into individual 500 mL pots containing the same sterilized substrate and inoculated with rhizobacteria. One week later, seedlings were either mechanically inoculated with BPMV or mock-inoculated, following the factorial design described in Table S1. All plants were maintained in an insect-free growth chamber at 25°C (day) and 23°C (night), with a 16:8 light/dark cycle and 70% relative humidity. We established BPMV infections by mechanical inoculation at the V1 growth stage, one week after bacterial inoculation. Half of the plants from each bacterial treatment were rub-inoculated with BPMV using a 0.1 M potassium phosphate buffer and carborundum powder; the remaining plants received virus-free buffer as a mock treatment. BPMV infection was verified at the end of the experiment by enzyme-linked immunosorbent assay (ELISA; Agdia).

Starting at the V1 growth stage, plants received 50 mL of a modified Hoagland’s nutrient solution three times per week (Pulido *et al*., 2024). Plants inoculated with *B. japonicum* alone or in combination with *D. acidovorans* received a nitrogen-free version of this solution, as nitrate supplementation inhibits nodule formation and biological nitrogen fixation (Ohyama *et al*., 2011; Carroll & Mathews, 2018). Uninoculated controls and *D. acidovorans*-only treatments received the standard solution including nitrogen. Although this introduces a nutritional difference between treatments, it reflects the conditions under which rhizobial symbiosis naturally occurs and is consistent with our prior work (Pulido *et al*., 2019, 2024) and comparable studies in rhizobial systems (Barros De Carvalho *et al*., 2013; Kontopoulou *et al*., 2015; Allito *et al*., 2021). We interpret our findings with this limitation in mind.

### Beetle colonies and larval performance assay

Colonies of the Mexican bean beetle (*Epilachna varivestis*), a major legume herbivore, were obtained from the Phillip Alampi Beneficial Insect Laboratory (New Jersey Department of Agriculture) and reared on *Phaseolus vulgaris* at 25°C under a 16:8 h light:dark photoperiod. Beetles were transferred to the experimental environment one day before assays. Larval performance was assessed in no-choice assays: ten newly hatched larvae were placed on each plant (n = 7 per treatment) and allowed to feed until the prepupal stage (approximately two weeks). Prepupae were collected, dried at 50°C for 48 h, and total dry mass recorded; mean weight per larva was calculated by dividing total dry mass by the number of surviving individuals. Mesh enclosures prevented larval movement between plants.

### Plant physical traits

Before grinding for metabolite extraction, leaf tissue was weighed and total leaf biomass recorded for each plant. Following aboveground harvest, roots were washed, nodules were collected, placed in labelled paper envelopes, and dried at 50°C for 48 h for dry mass determination. Since nodules form exclusively in *B. japonicum*-inoculated plants, control and *D. acidovorans*-treated plants were also examined to verify the absence of cross-contamination. Virus symptom severity was assessed on an ordinal scale of 1–5 at the end of the experiment.

### Metabolomic extraction and analysis

Detailed protocols for metabolite extraction, data acquisition, processing, and compound identification have been described previously (Pulido *et al*., 2024). Briefly, leaf tissue was collected after harvest, flash-frozen in liquid nitrogen, and lyophilized (n = 5 per treatment). Dried tissue (10 ± 0.6 mg) was homogenized and subjected to a multi-phase extraction protocol to isolate polar, non-polar, and secondary metabolites. Metabolite profiling was performed using gas chromatography-mass spectrometry (GC-MS) and ultra-performance liquid chromatography-tandem mass spectrometry (UPLC-MS/MS) in both positive and negative ion modes.

Raw GC-MS files were processed in Agilent Quantitative MassHunter, with peak detection and deconvolution followed by putative compound identification against the NIST 2014 spectral library. LC-MS/MS raw files were processed in MZmine3 (Schmid *et al*., 2023) with intensity thresholds set at 1 × 10³ (MS1) and 1 × 10² (MS/MS). Extracted ion chromatograms were built using the ADAP chromatogram builder (minimum peak intensity 1 × 10²; mass window 20 ppm), and peaks were deconvoluted using the local minimum resolver (chromatographic threshold 90%; minimum peak height 5 × 10³). Feature alignment used the join aligner function (m/z tolerance 5 ppm; retention time tolerance 0.1 min) after filtering for ¹³C isotopes. Putative identification was based on spectral similarity with the MoNA library (Vaniya *et al*., 2019) using a spectral m/z tolerance of 20 ppm; MS2 spectra were further processed for in-silico annotation and compound classification via SIRIUS (Dührkop *et al*., 2019) and NPClassifier (Kim *et al*., 2021). Feature tables from each extraction phase were normalized using phase-specific internal standards and initial sample weights before merging with GC-MS data. Compound names were assigned based on identifiers listed in Dataset S4.

### Data analysis

All statistical analyses were conducted in R v.4.4.0 (R. Core Team, 2016). Mean larval weight was analysed using a generalized linear mixed model (GLMM) with soybean variety, rhizobacteria treatment, and virus infection as fixed effects. Larval survival was modelled with a separate GLMM using the glmmTMB package (Brooks *et al*., 2017), with the number of surviving larvae out of 10 individuals as the response variable, a binomial error distribution, and a logit link function. Predicted survival probabilities and standard errors were extracted from the fitted model. Pairwise odds ratios (ORs) and 95% confidence intervals (CIs) for key contrasts—including RR vs NonRR within each rhizobacteria × virus combination—were computed using estimated marginal means (emmeans), where an OR > 1 indicates higher survival in the first group.

Shoot, root, and nodule biomass were each analyzed independently using GLMMs with Gaussian error distributions (*glmmTMB*), with rhizobacteria treatment, virus infection, soybean variety, and their three-way interactions as fixed effects. Estimated marginal means for all rhizobacteria × virus × variety combinations were computed using *emmeans*, and pairwise RR vs NonRR contrasts were extracted to isolate variety-specific effects under each treatment combination. The relationship between nodule biomass and shoot biomass was further examined by regression analysis, conducted separately for each soybean variety across rhizobacteria × virus treatment combinations, to evaluate whether nodulation influenced overall plant growth under varying microbial contexts.

Symptom severity (ordinal scale 1–5) was analysed using a cumulative link model (CLM) with a logit link function, implemented with the clm function from the ordinal package (Christensen, 2019). Rhizobacteria treatment and soybean variety were included as fixed effects, with virus symptom severity treated as an ordered factor.

To accommodate the high dimensionality and heterogeneous structure of the combined metabolite dataset, we first applied Discriminant Analysis of Principal Components (DAPC; Jombart *et al*., 2010) to visualize global separation among treatment groups. We then employed Weighted Gene Co-expression Network Analysis—applied here to metabolite co-abundance data (WGCNA; Langfelder & Horvath, 2008)—to identify modules of co-abundant metabolites (soft-thresholding power β = 12). Module eigengenes (MEs) were used to assess correlations between metabolite modules and experimental factors, providing an integrated view of treatment-associated metabolome reorganization.

## Results

### Effects of RR gene presence and microbe colonization on larval performance

#### Weight gain in larvae

BPMV infection and rhizobacterial inoculation both enhanced larval weight gain, while the RR transgene had no significant main effect on this measure. Specifically, larvae gained significantly more weight on BPMV-infected plants than on uninfected ones (Fig. 1A, Table S2, p = 0.035). Single inoculation with *B. japonicum* (Bj) also increased larval weight (Fig. 1A, Table S2, p = 0.038), and this effect was further enhanced under dual inoculation with *D. acidovorans* (BjDa) (Fig. 1A, Table S2, p = 0.013). In contrast, single inoculation with Da alone showed a marginal negative effect on larval weight (Fig. 1A, Table S2, p = 0.081).

**Figure 1.**
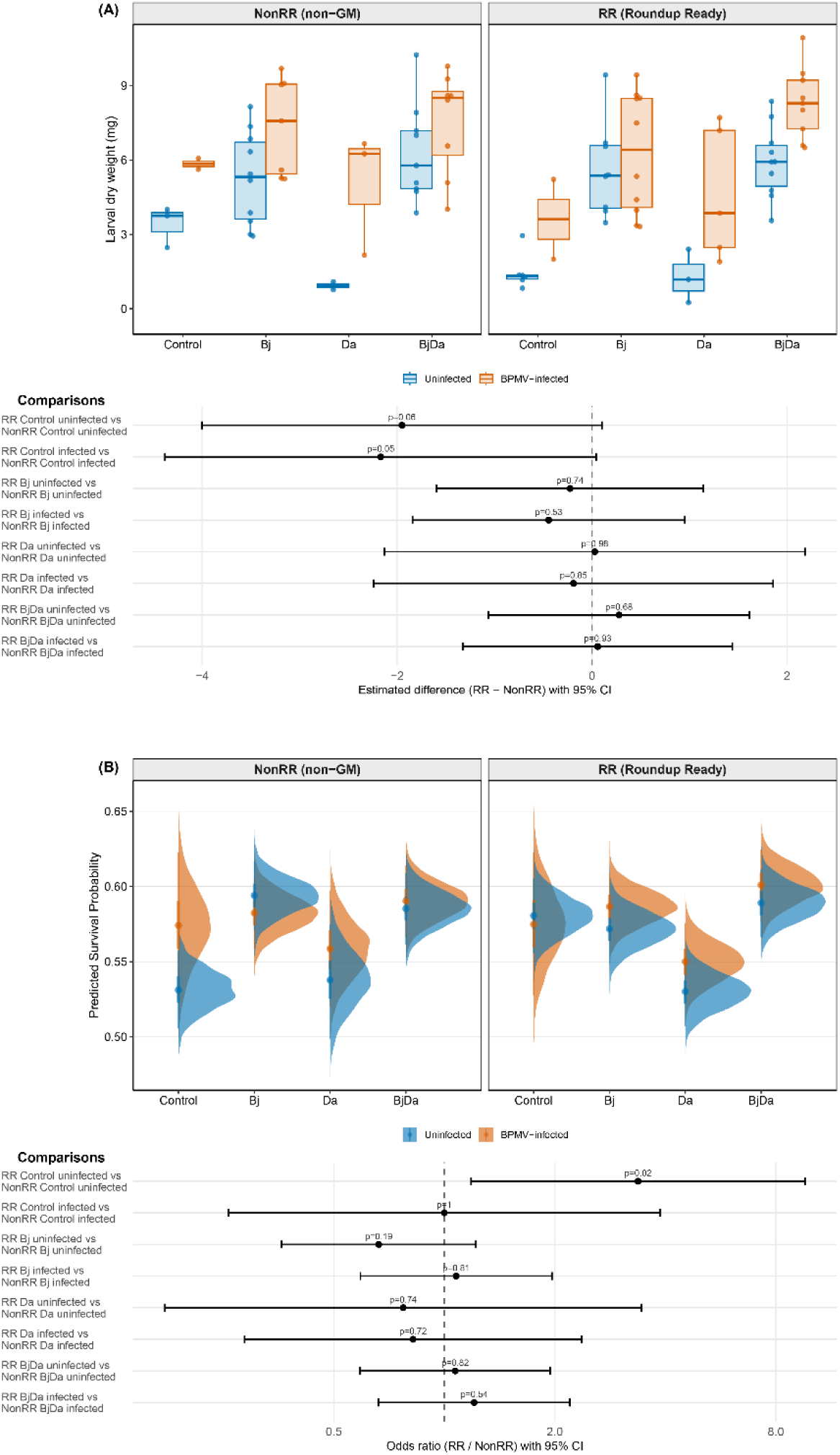
Effects of rhizobacteria inoculation, BPMV infection, and soybean variety on larval performance of Epilachna varivestis. (A) Larval dry weight at the prepupal stage. **Top (Response plot)**: Raw data distribution with treatment means (dots) and 95% CIs (error bars). Rhizobacteria treatment is shown on the x-axis; color indicates virus infection status (blue: uninfected; red: BPMV-infected). **Bottom(Effect plot):** Pairwise contrasts between RR and NonRR plants within each rhizobacteria × virus combination (effect sizes with 95% CIs from GLMM); positive values indicate higher weight on RR plants. (B) Larval survival probability (proportion surviving to prepupal stage out of 10 individuals per plant). **Top (Response plot)**: GLMM-predicted survival probabilities with 95% CIs; color as in (A). **Bottom (Effect plot)**: Pairwise contrasts between RR and NonRR plants within each rhizobacteria × virus combination, expressed as odds ratios (95% CIs from GLMM); values greater than 1 indicate higher survival odds on RR plants. NonRR: near-isogenic non-GM soybean; RR: Roundup Ready soybean; Bj: Bradyrhizobium japonicum-inoculated; Da: Delftia acidovorans-inoculated; BjDa: co-inoculated with Bj and Da.

The Roundup Ready (RR) variety did not significantly impact larval weight (Fig. 1A, Table S2, p = 0.059). However, pairwise comparisons revealed a marginal reduction in larval weight in uninfected and not inoculated RR plants (Fig. 1A, Table S3, p = 0.06), an effect that was further decreased under BPMV infection (Fig. 1A, Table S3, p = 0.05). Notably, the positive effects of rhizobacteria on larval performance were consistent across both soybean varieties, as no significant interactions between rhizobacteria and variety were detected. Full model results are reported in Table S3.

#### Larval survival

Larval survival was significantly influenced by rhizobacterial inoculation and soybean variety, but not by virus infection alone (Fig. 1B, Table S4). Inoculation with *B. japonicum* increased survival odds by 1.5-fold relative to controls (Fig. 1B, Table S4, p = 0.005), and dual inoculation (BjDa) produced a similar effect (Fig. 1B, Table S4, p = 0.013); single *D. acidovorans* inoculation had no significant effect (Fig. 1B, Table S4, p = 0.8).

The Roundup Ready (RR) variety also significantly increased larval survival, with larvae being 1.2 times more likely to survive on RR plants than on the non-GM line (NonRR) (Fig. 1B, Table S4, p = 0.023). However, pairwise contrasts revealed that this effect was only significant for control uninfected plants, where RR plants increased survival odds by 3.37 times compared to NonRR controls (Fig. 1B, Table S5, p = 0.02). Interestingly, the positive effect of rhizobacteria on survival was reduced when combined with the RR variety, particularly in Bj-inoculated plants (Fig. 1B, Table S4, p = 0.009). Additionally, no significant interactions were detected between rhizobacteria and virus infection, nor between virus infection and variety, suggesting that virus infection alone does not alter larval survival, but rather the interaction between rhizobacteria and the plant genetic background plays a key role in shaping survival outcomes. Full model results are reported in Table S4.

### Plant physical traits

Virus infection significantly reduced shoot biomass across all treatments, regardless of soybean variety or rhizobacteria inoculation (Fig. 2A, Table S6, p < 0.0001). In contrast, dual inoculation with *B. japonicum* and *D. acidovorans* (BjDa) enhanced shoot biomass (Fig. 2A, Table S6, p = 0.049), while single inoculation with Da resulted in a reduction (Fig. 2A, Table S6, p = 0.006). The Roundup Ready (RR) variety produced greater shoot biomass than the nonRR line, but only in Bj-inoculated and uninfected plants (Fig. 2A, Table S7, p = 0.01). No significant synergistic interactions were detected between variety, rhizobacteria, and virus treatments.

**Figure 2.**
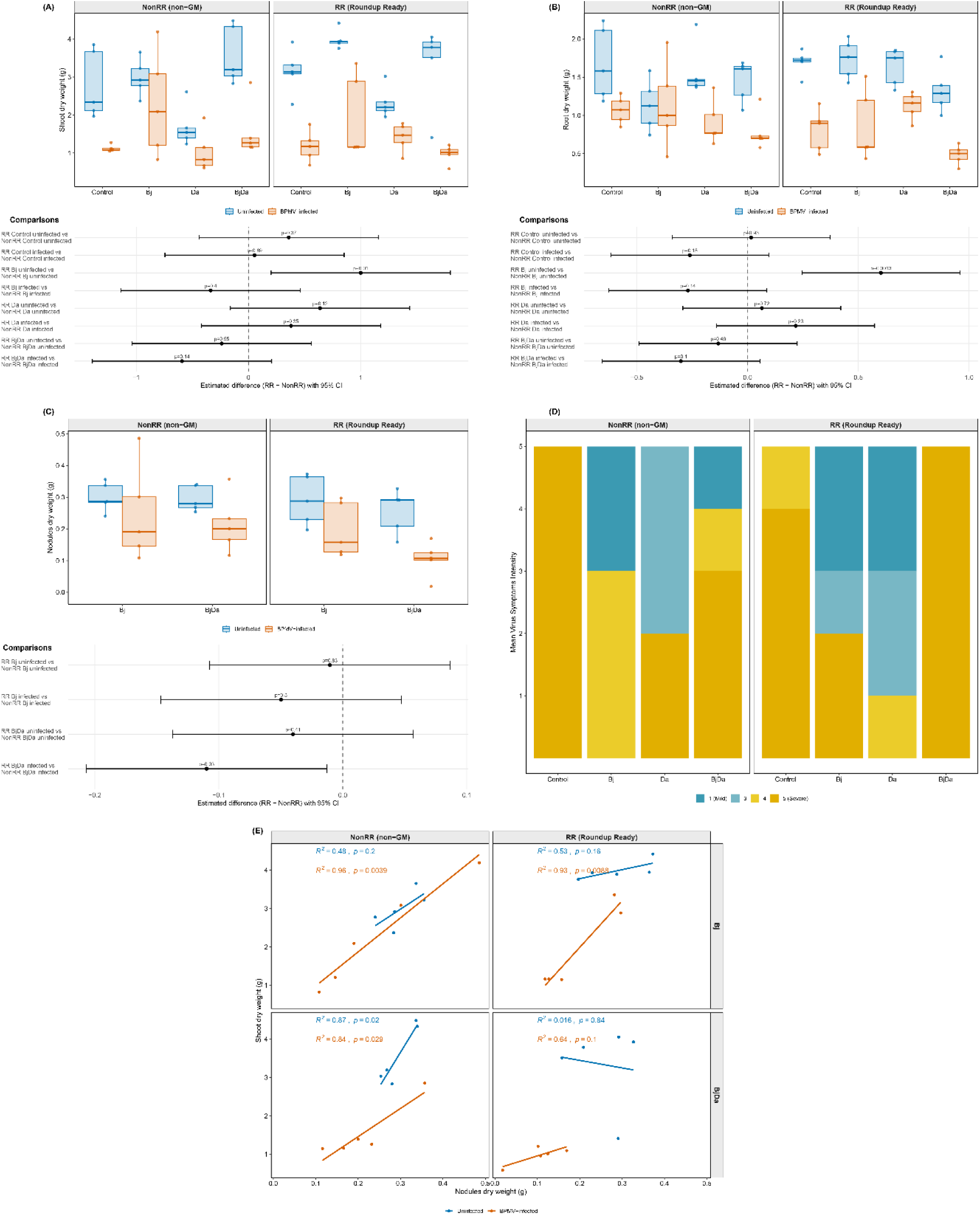
Effects of rhizobacteria inoculation, BPMV infection, and soybean variety on plant physical traits. (A) Shoot biomass, (B) root biomass, and (C) nodule biomass. **Top panels**: Raw data distribution per treatment group. Large dots represent group means; error bars indicate 95% CIs. Rhizobacteria treatment is shown on the x-axis; color indicates virus infection status (blue: uninfected; red: BPMV-infected). Nodule biomass is shown only for B. japonicum-inoculated treatments, as nodules do not form in uninoculated or D. acidovorans-only plants. **Bottom panels**: Pairwise contrasts between RR and NonRR plants within each rhizobacteria × virus combination (effect sizes with 95% CIs from GLMM); positive values indicate greater biomass in RR plants. (D) Virus symptom severity across rhizobacteria treatments and soybean varieties. Stacked bars show the number of plants in each symptom severity category (1 = no visible symptoms; 5 = severe systemic mosaic). Both B. japonicum and D. acidovorans inoculation significantly reduced symptom severity regardless of plant genotype. (E) Relationship between nodule biomass and shoot biomass in B. japonicum-inoculated (Bj) and co-inoculated (BjDa) plants, shown separately for each soybean variety and virus infection status. Slopes (β) represent the change in shoot biomass (g) per 1 g increase in nodule biomass from a linear mixed-effects model. A significant positive relationship was detected only in BPMV-infected plants, suggesting that nodulation partially compensates for virus-induced biomass losses. NonRR: near-isogenic non-GM soybean; RR: Roundup Ready soybean; Bj: Bradyrhizobium japonicum-inoculated; Da: Delftia acidovorans-inoculated; BjDa: co-inoculated with Bj and Da.

Similarly, virus infection reduced root biomass across all conditions (Fig. 2B, Table S8, p < 0.001). While single inoculation with *B. japonicum* decreased root biomass, virus infection counteracted this effect, restoring root biomass levels in *B. japonicum*-inoculated plants (Fig. 2B, Table S8, p = 0.016). Additionally, an interaction effect was observed in RR plants, where *B. japonicum* inoculation increased root biomass, but only in virus-free conditions (Fig. 2B, Table S9, p = 0.020).

Nodulation was not significantly affected by treatment interactions (Fig. 2C, Table S10). However, pairwise comparisons revealed that RR plants exhibited reduced nodule biomass when dual-inoculated and infected with BPMV (Fig. 2C, Table S11, p = 0.03).

Both *B. japonicum* (Fig. 2D, Table S12, p = 0.003) and *D. acidovorans* (Figure Fig. 2D, Table S12, p = 0.004) significantly reduced virus symptoms, but plant variety had no effect on symptom severity.

A linear regression model assessing shoot biomass as a function of nodule biomass revealed that virus infection significantly strengthened this relationship (Fig. 2E, Table S13, p = 0.023). In infected plants, shoot biomass increased sharply with nodule biomass, both in NonRR plants and RR plants under single and dual inoculation (Bj and BjDa) (Fig. 2E, Table S14, p < 0.0001). In contrast, this relationship was not significant in uninfected plants, where nodule biomass had minimal influence on shoot biomass. Neither dual inoculation nor plant variety significantly altered this relationship, suggesting that the compensatory role of nodules in shoot biomass production under viral stress is independent of these factors.

### Effects on plant metabolomics

To characterize treatment-driven metabolic shifts, we integrated data from four extraction phases — LC-MS/MS in both negative and positive ion modes and GC-MS water and chloroform fractions — yielding approximately 1,529 metabolite features. Of these, 217 were putatively matched to public spectral libraries, and 336 were annotated with chemical class information using NPClassifier (Kim *et al*., 2021). Annotated metabolites spanned all major pathways—including amino acids, carbohydrates, fatty acids, flavonoids, isoflavonoids, shikimate–phenylpropanoids, terpenoids, and polyketides—providing a broad basis for assessing soybean metabolic responses.

To explore treatment effects, we first applied WGCNA to organize metabolites into co-abundance modules and tested their associations with treatments (Fig. 4A–C). Between-genotype comparisons (RR vs. NonRR) highlighted modules with divergent responses to rhizobacterial inoculation and viral infection (Fig. 4B; Dataset S1). In parallel, within-genotype baseline contrasts revealed substantial shifts in compound abundance in response to microbial inoculation and infection (Dataset S2), with the distribution of differentially accumulated metabolites (DAMs) per pathway class summarized in Figs. S1 (NonRR) and S2 (RR). In NonRR plants, rhizobacterial inoculation consistently elevated metabolites across amino acid, carbohydrate, and flavonoid pathways, with the broadest response occurring under B. japonicum single inoculation and dual (Bj+Da) inoculation; BPMV infection further expanded the number of significantly altered metabolites, particularly reducing shikimate–phenylpropanoid and fatty acid compounds in several treatments (Fig. S1). In RR plants, the overall pattern was qualitatively similar but differed in magnitude and pathway balance: rhizobacterial treatments elevated isoflavonoids and amino acid-related metabolites more selectively, while suppressing a broader set of lipid-related and terpenoid compounds, an asymmetry that intensified under viral co-infection (Fig. S2). These genotype-specific differences in the breadth and directionality of pathway responses form the metabolic substrate for the herbivore performance outcomes described below. This complementary framework allowed us to distinguish genotype-specific patterns from inoculation-driven effects that were consistent across genetic backgrounds.

**Figure 3.**
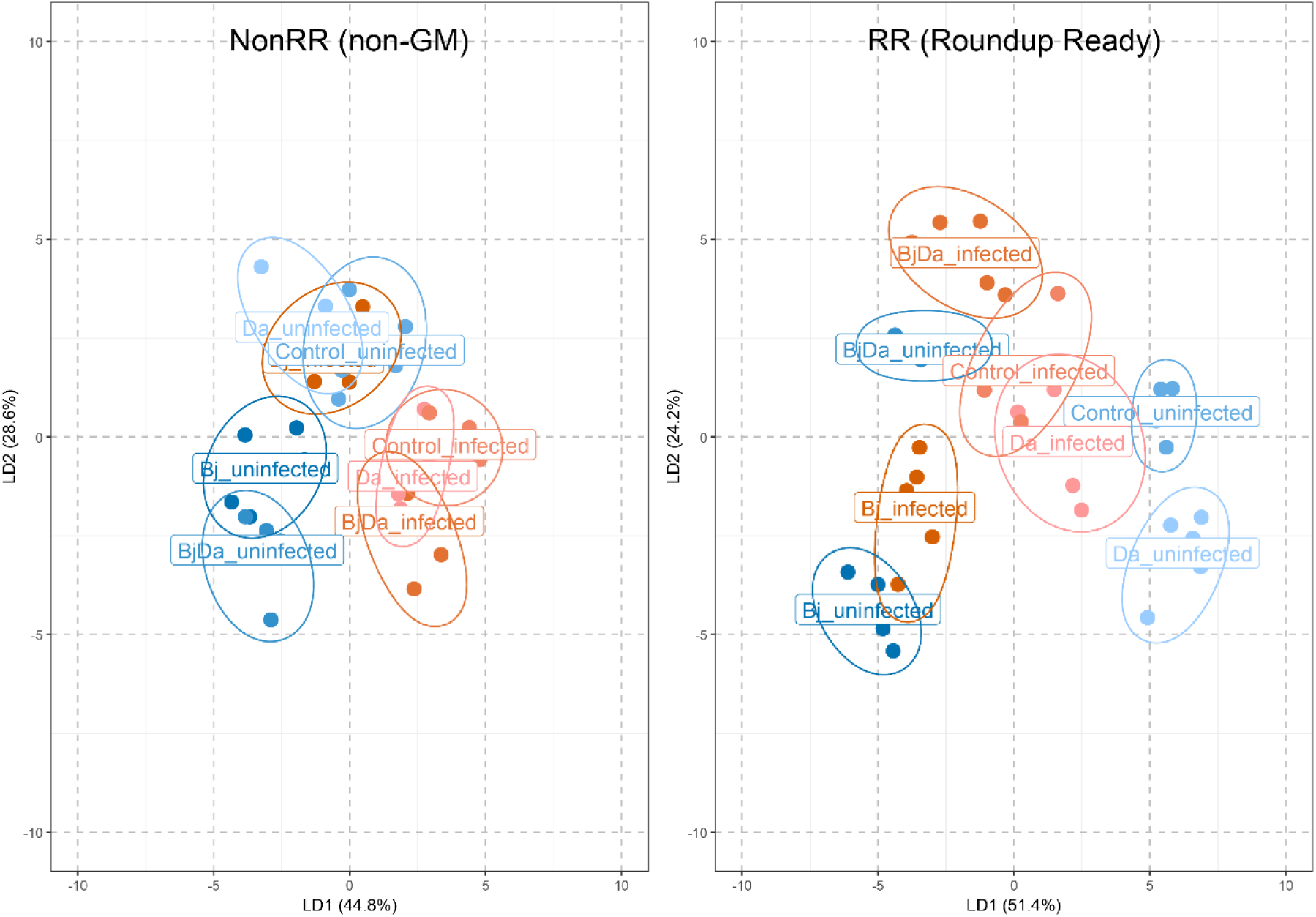
Metabolic separation among soybean treatment groups revealed by Discriminant Analysis of Principal Components (DAPC). Each panel shows DAPC scores for NonRR (left) and RR (right) plants, plotted along Discriminant Functions 1 and 2. Each point represents an individual plant sample, color-coded by rhizobacteria × virus treatment. Ellipses represent 95% inertia ellipses for each treatment group. Virus infection status is the primary driver of metabolic separation in both genotypes, with infected and uninfected groups forming distinct clusters. Rhizobacterial inoculation, particularly B. japonicum single and dual inoculation (Bj, BjDa), amplifies this separation, with BjDa treatments diverging furthest from controls. Separation among treatment groups is slightly more compact in RR than in NonRR plants, consistent with genotype-dependent differences in metabolic responsiveness under multi-species stress. NonRR: near-isogenic non-GM soybean; RR: Roundup Ready soybean; Bj: Bradyrhizobium japonicum-inoculated; Da: Delftia acidovorans-inoculated; BjDa: co-inoculated with Bj and Da.

**Figure 4.**
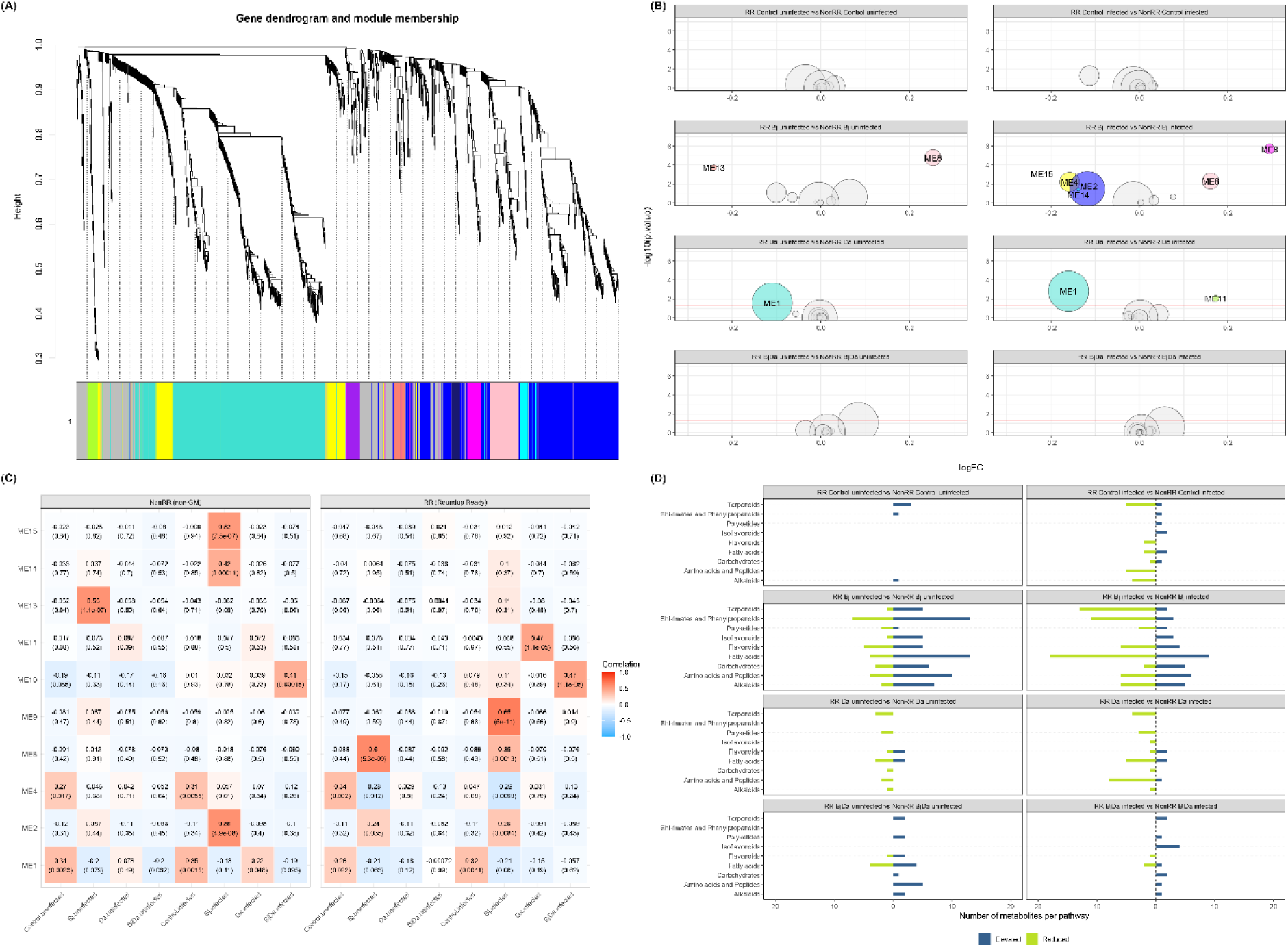
Weighted co-expression network analysis (WGCNA) of the soybean metabolome under rhizobacterial inoculation and BPMV infection in RR and NonRR varieties. (A) Hierarchical clustering dendrogram of metabolite features based on topological overlap, constructed jointly from RR and NonRR plants. Branch height reflects dissimilarity between modules; each module is assigned a unique color identifier by WGCNA. Ten biologically interpretable modules were identified (excluding the gray module of unassigned low-connectivity features, n = 245). (B) Differential module abundance between RR and NonRR plants across all rhizobacteria × virus treatment combinations, based on linear model contrasts. Each bubble represents a module showing significant differential abundance in at least one treatment contrast; bubble size indicates the number of significantly differentially accumulated metabolites (DAMs) within that module, and bubble position on the x-axis reflects the direction and magnitude of the log-fold change (RR relative to NonRR). Modules shifting upward in RR plants are shown above zero; those reduced in RR plants are shown below. Treatments are shown on the y-axis. (C) Module-trait correlation heatmap showing associations between module eigengenes and treatment combinations across both genotypes. Cell color reflects the strength and direction of the Pearson correlation; significant associations (p < 0.05) are indicated. This panel captures treatment-driven co-expression patterns shared across genotypes, complementing the genotype-specific contrasts shown in (B). (D) Number of differentially accumulated metabolites (DAMs) per metabolic compound class across RR vs. NonRR contrasts for each rhizobacteria × virus treatment. Bars show the count of DAMs per class; upward bars indicate metabolites more abundant in RR plants and downward bars indicate metabolites more abundant in NonRR plants. This panel illustrates which compound classes drive genotype-specific metabolic divergence under each treatment combination. NonRR: near-isogenic non-GM soybean; RR: Roundup Ready soybean; Bj: Bradyrhizobium japonicum-inoculated; Da: Delftia acidovorans-inoculated; BjDa: co-inoculated with Bj and Da.

#### Metabolomic Profiling Reveals Distinct Treatment-Driven Separations in NonRR and RR Soybeans

Discriminant Analysis of Principal Components (DAPC) (Fig. 3) revealed structured metabolic separation among soybean treatments, driven mainly by virus infection status. In the NonRR variety, infected and uninfected groups form distinctly separated clusters, especially in the *B. japonicum* (Bj) and dual inoculation (BjDa) treatments. The dual inoculation group diverges furthest from controls, indicating a potential synergistic interaction between *B. japonicum* and *Delftia acidovorans* (Da). In the RR variety, a similar virus-driven separation is observed, though slightly more compact and less spread than in NonRR. Notably, the Bj and BjDa treatments again show enhanced separation, suggesting that rhizobacteria inoculation amplifies virus-induced metabolic divergence, particularly in Roundup Ready soybeans. The Da-only treatments show intermediate positioning, potentially indicating a more subtle effect. Together, these results indicate that both infection status and rhizobacterial inoculation, especially in combination, shape the soybean metabolomic profile in a genotype-dependent manner.

#### WGCNA: Module-Trait Correlations and the Influence of Roundup Ready Soybean

WGCNA revealed no major differences in co-correlation patterns between soybean varieties (Fig. 4). The network grouped metabolites into ten distinct modules as visualized in the gene dendrogram (Fig. 4A). Compounds with similar expression profiles were grouped into modules, each represented by a unique color. The largest modules were turquoise (ME1, n = 488) and blue (ME2, n = 376), while midnightblue (ME15, n = 24) was the smallest. A set of 245 metabolite features was not assigned to any module and were grouped into the gray category, typically indicating low connectivity. These modules formed the basis for downstream correlation and enrichment analyses.

#### RR vs NonRR in non-inoculated (Control) plants and plants infected with BPMV

In the absence of microbial inoculation or viral infection, no WGCNA modules showed significant changes in abundance, and module-trait correlations were weak (Fig 4B, 4C), suggesting minimal baseline divergence in global metabolomic structure between genotypes. However, compound-level differential abundance analysis revealed that even under baseline conditions, RR plants exhibited increased levels of a small set of metabolites. Notably, several compounds assigned to ME1—including the photosynthetic pigment pheophytin a, and triterpenoid-related compounds—showed significantly higher abundance in RR plants (Fig. 4D, Dataset S1). These findings suggest that the glyphosate-tolerant trait alone introduces minimal disruption to metabolic organization in the absence of biotic stress, reinforcing the principle of substantial equivalence and consistent with prior work suggesting near-isogenic similarity under unstressed conditions (Barros *et al*., 2010).

Upon BPMV infection, metabolic distinctions between RR and nonRR plants became more pronounced. While no WGCNA modules reached significance in module-level limma analysis, compound-level comparisons revealed a broader range of differentially abundant metabolites across multiple pathways, particularly in modules ME1 and ME4. Several metabolites associated with multiple modules—such as isorhamnetin-3-galactoside, Beta-Sitosterol and pinnatasterone—showed significant reduction in BPMV-infected RR plants relative to BPMV-infected nonRR plants (Fig. 4D, Dataset S1). These compounds span flavonoid, fatty acids, amino acids and alkaloids pathways, many of which are involved in viral defense and oxidative signaling. In general, we observed that viral infection introduces greater metabolic divergence between RR and nonRR plants, even in the absence of microbial inoculation.

#### RR vs NonRR in B. japonicum inoculated plants without and with BPMV infection

In uninfected plants inoculated with *B. japonicum* (Bj), RR plants showed strong positive correlation with ME8 (r = 0.6, p < 0.0001, Fig. 4C), a module enriched in amino acids (e.g., L-phenylalanine), nucleosides and flavonoids. This association aligns with compound-level results (Dataset S1), where multiple isoflavones (e.g., malonyldaidzin, formononetin derivatives), flavonols (e.g., isorhamnetin, kaempferol 3-rhamnoside), and saccharides (e.g., D-glucose) accumulated at higher levels in RR plants colonized with Bj and lacking BPMV infection. Conversely, ME13 was significantly reduced in uninfected RR plants colonized with Bj relative to uninfected Bj-colonized nonRR plants (logFC = –0.24, p < 0.001, Fig. 4B), and compound-level data revealed consistent depletion of fatty acids and triterpenoids.

Upon BPMV infection, metabolic differences between Bj-colonized RR and nonRR plants intensified. Infected RR plants with Bj colonization increased ME9 (logFC = 0.65, p < 0.0001, Fig. 4B) and ME8 (logFC = 0.35, p = 0.0013, Fig. 4B), while decreasing ME2, ME4, ME14, and ME15 relative to infected nonRR plants with Bj colonization (Fig. 4B). Compound-level data showed broad suppression of flavonoids, phenylpropanoids, fatty acids, and alkaloids—key classes in generalized defense—alongside selective accumulation of isoflavonoids (e.g., ononin, formononetin derivatives), amino acids, and nucleosides (Dataset S1). The overall pattern suggests a reorientation toward nitrogen assimilation and specialized secondary metabolism in BPMV-infected RR plants colonized by Bj, relative to infected Bj+ nonRR plants, potentially at the cost of broad defense pathways. Within ME2 and ME4, responses were more heterogeneous: while some phenylpropanoids (e.g., ferulic acid, caffeic acid derivatives) and flavones (e.g., chrysin) were suppressed in RR plants, others such as fatty acids were significantly elevated (Fig. 4E, Dataset S1). Despite the overall reduction of compounds in these pathways, several isoflavonoids, including ononin and formononetin malonylglucoside, were significantly elevated in RR plants (Dataset S1).

#### RR vs NonRR in D. acidovorans inoculated plants without and with BPMV infection

In the absence of viral infection, metabolic differences between RR and nonRR plants inoculated with *D. acidovorans* (Da) were relatively subtle. At the module level, only ME1 showed a significant reduction in module activity in RR plants inoculated with Da (*logFC* = – 0.11, *p* = 0.028, Fig. 4B). Compound-level analysis revealed a modest suppression of several amino acids, fatty acids, and triterpenoid saponins, along with decreased levels of the terpenoid Soyasaponin III. Some flavonoids, including luteolin and quercetin, were elevated, while others (e.g., apigenin) were reduced, indicating a non-uniform response across subclasses (Dataset S1).

Following BPMV infection, RR plants inoculated with *D. acidovorans* exhibited similar trends to the infected vs. uninfected condition but with enhanced suppression of key metabolic groups. ME1 was again significantly reduced (*logFC* = –0.16, *p* = 0.0016, Fig. 4B), while ME11 showed moderate upregulation (*logFC* = 0.17, *p* = 0.009, Fig. 4B; r = 0.47, p < 0.0001, Fig. 4C), consistent with lipid remodeling. At the compound level, a broader decline in amino acids, fatty acid derivatives, and various triterpenoids (e.g., soyasaponin III, soyasapogenol B) was observed for BPMV-infected RR plants inoculated with *D. acidovorans* relative to BPMV-infected nonRR plants with *D. acidovorans*. Isoflavonoids and some flavonols (e.g., quercetin glucosides) remained elevated, while others (e.g., narcissin) declined, reinforcing a selective modulation of secondary metabolism. BPMV-infected NonRR plants did not display any significant module associations under *D. acidovorans* treatments.

#### RR vs NonRR in dual rhizobacteria inoculation (BjDa) without and with BPMV infection

RR plants co-inoculated with *B. japonicum* and *D. acidovorans* (BjDa) exhibited a strong positive correlation with ME10 (r = 0.470, p < 0.0001, Fig. 4C), a module dominated by isoflavonoids and saccharides, suggesting induction of shikimate-phenylpropanoid defenses.

In uninfected plants, dual rhizobacterial inoculation (BjDa) elicited fewer metabolic changes in RR plants compared to the single *B. japonicum* treatment. No WGCNA modules were significantly altered, and only a limited set of compounds showed significant differential abundance, including cyclic and linear peptides (e.g., hirsutatin A), several flavones (e.g., luteolin, 7,4’-dihydroxyflavone), and saponins (e.g., soyasaponin I, III). In contrast, certain fatty acids (e.g., palmitic acid, 2,3-butanediol) and flavonols (e.g., quercetin glycosides) were suppressed (Fig. 4D, Dataset S1).

BPMV infection moderately amplified these metabolic shifts, particularly enhancing isoflavonoid accumulation (e.g., biochanin B, isoflavone conjugates in ME1 and ME10).

Saponins such as **soyasaponin I** and **III** and their aglycones (soyasapogenol B) also increased in RR plants, consistent with an increased defense response. However, suppression of quercetin glycosides, diacylglycerols, and dicarboxylic acids (e.g., methylmalonic acid) indicates possible trade-offs in primary metabolism and membrane remodeling. Despite these compound-level changes, no WGCNA modules showed significant shifts.

## Discussion

RR soybean plants exhibited distinct patterns of metabolic regulation under multi-species biotic stress, notably through selective shifts in lipid metabolism and shikimate-phenylpropanoid pathways, while remaining metabolically indistinguishable from their non-GM counterparts under baseline conditions. These metabolomic changes mirrored ecological outcomes: BPMV infection generally increased Mexican bean beetle (*Epilachna varivestis*) larval weight gain across genotypes, while rhizobacterial inoculation promoted shoot growth. However, the underlying metabolic changes varied depending on the rhizobacteria inoculated to soybean roots. Elevated isoflavonoid accumulation in some infected RR plants (especially under BjDa) likely enhanced antimicrobial defenses without directly deterring herbivores, whereas the suppression of amino acids and fatty acids under *D. acidovorans* inoculation may have reduced the nutritional quality of the plants, decreasing herbivore growth as well. Our results indicate that the modified EPSPS gene has minimal impact on plant metabolites in isolation, but measurably alters how plants integrate and respond to concurrent microbial and viral signals. These differences are not detectable under simplified experimental conditions — they only surface when plants are challenged by the kind of multi-species biotic context that characterizes real agricultural environments. This context-dependence is itself an ecologically important finding: it means that transgene effects on tritrophic dynamics can be invisible in single-species assays yet consequential at the community level.

### Genotype-specific rewiring of metabolic responses under microbial and viral challenge

In the absence of microbial inoculation or viral infection, metabolomic differences between RR and NonRR soybeans were minimal. Neither WGCNA module structure nor trait correlations showed significant genotype effects, supporting the principle of substantial equivalence under unstressed conditions (Barros *et al*., 2010). Nevertheless, compound-level analysis revealed subtle but consistent differences: RR plants exhibited elevated abundance of specific triterpenoids and pigments, such as pheophytin a. These findings suggest that the transgenic event has a subtle effect on some compounds without disruption of primary metabolic pathways. Upon BPMV infection, divergences became more apparent. RR plants showed reductions in key defense-related metabolites, including isorhamnetin-3-galactoside and fatty acid conjugates. Although no single module dominated the infection response, the cumulative suppression of flavonoids, fatty acids, and amino acids in RR plants points toward weakened chemical defenses enhancing larval survival, while the concurrent depletion of amino acids and fatty acids conjugates limits nutritional quality, explaining the higher survival but reduced weight gain. This aligns with studies showing that specialist herbivores tolerate defensive metabolites but suffer growth penalties when host nutritional capacity is compromised (Rothwell & Holeski, 2020).

*Bradyrhizobium japonicum* inoculation elicited stronger and more complex metabolic shifts than *D. acidovorans* treatments. In uninfected conditions, RR plants showed increased module activity of **ME8**—rich in amino acids and nucleosides—and increased levels of several isoflavonoids and flavonols. These changes likely reflect enhanced nitrogen assimilation and secondary metabolism, typical of successful rhizobial symbiosis. Meanwhile, fatty acids and triterpenoids decreased, suggesting trade-offs in resource allocation. Upon BPMV infection, RR plants diverged further from their non-GM counterparts. They exhibited a shift toward ME9 (glycerolipids and isoflavonoid conjugates) and suppression of defense modules (ME2, ME4, ME15). Notably, the reduction of ME15—a module rich in antimicrobial terpenoids and fatty acid derivatives such as sesquiterpenoids and diterpenoids—in infected RR plants with Bj colonization highlights a potential compromise in basal defense capacity.

These patterns align with previous reports of suppressed secondary metabolism in genetically modified maize non-herbicidal conditions (Mesnage *et al*., 2016), suggesting that such trade-offs may become more apparent under multifactorial stress conditions. Taken together, these findings indicate that under *B. japonicum* colonization and viral challenge, RR plants divert metabolic investment from broad-spectrum defenses toward a narrower, more specialized set of stress-responsive compounds. This reallocation — absent under single-species conditions — illustrates how the functional consequences of a transgene can depend entirely on the complexity of the biotic environment in which it is expressed.

Inoculation with *Delftia acidovorans* alone elicited less extensive metabolic changes than with *B. japonicum*. Uninfected RR plants displayed a suppression of **ME1**, alongside slight elevations in certain flavonoids (e.g., luteolin, quercetin). BPMV infection further suppressed **ME1** and induced a strong association with **ME11** in RR plants—a module enriched in fatty acids and polyketide derivatives, including 1-octacosanol. These compounds are frequently associated with structural defenses and membrane-based signaling (Kachroo & Kachroo, 2009), suggesting that RR plants prioritize lipid remodeling under combined microbial and viral stress. The reduced abundance of metabolites in **ME1** points to a metabolic reallocation under dual biotic pressure. **ME1** comprises a chemically diverse set of metabolites, including photosynthetic pigments (pheophytin a), carotenoids, flavonol glycosides (kaempferol-3-O-glucoside), membrane lipids (1-18:3-lysoPC, monolinolenin), and defense compounds such as D-pinitol, a sugar alcohol that has been observed to increase in response to abiotic stress (Dumschott *et al*., 2019) and a natural deterrent against insect herbivores (Liu *et al*., 2022).

The reduction of these compounds suggests a dampening of pathways involved in photoprotection, membrane stability, and osmotic balance, particularly under combined microbial colonization and viral infection. Compared to *B. japonicum* treatments, the overall magnitude and complexity of metabolic responses were lower, suggesting that colonization by a generalist rhizobacterium induces more limited metabolic reprogramming than colonization by a specialist symbiont. Consistent across both infected and uninfected states, RR plants showed broad reductions in amino acids, fatty acids, and saponins—traits likely important for structural integrity and herbivore resistance. The persistence of elevated flavonoid levels suggests some maintenance of chemical defenses. Overall, these findings align with the idea that specialized symbionts like *B. japonicum* engage more deeply with host metabolism than generalist symbionts (Oldroyd, 2013; Scott *et al*., 2021).

Dual inoculation of *B. japonicum* and *D. acidovorans* in RR plants (both uninfected and BPMV-infected) strongly correlated with **ME10**, a module dominated by shikimate–phenylpropanoid pathway metabolites, including isoflavonoids such as coumestrol and daidzein. These compounds are central to legume immunity and symbiotic signaling (Subramanian *et al*., 2007; Weston & Mathesius, 2013), and their selective accumulation in RR plants suggests that the glyphosate-tolerant trait may shift the balance of metabolic priorities toward specialized secondary metabolites when inoculated with specialist and generalist rhizobacteria. BPMV infection further increased isoflavonoids (e.g., biochanin B, coumestrol conjugates) and saponins, consistent with enhanced specialized defenses. The suppression of quercetin glycosides (defensive alkaloids) and membrane lipids suggests trade-offs where the plant may prioritize specialized defenses at the expense of maintaining foundational biochemical functions like membrane stability and photoprotection, processes known to be supported by flavonoids like quercetin under abiotic stress (Nakabayashi *et al*., 2014). These observed shifts in the shikimate–phenylpropanoid pathway are of particular relevance given the nature of the RR modification. The CP4 EPSPS enzyme confers glyphosate tolerance by maintaining the function of the shikimate pathway for aromatic amino acid synthesis even under herbicide application. Our results suggest that this modified enzymatic function may also influence the baseline flow of carbon through this pathway and its connected secondary metabolic branches, even in the absence of glyphosate. The selective accumulation of specific isoflavonoids (e.g., coumestrol, daidzein) in RR plants under dual inoculation, rather than a broad-scale activation, points to a complex interaction between the CP4 EPSPS gene and microbial signaling networks that regulate plant defense investment. This indicates that the transgene’s effects are not neutral in ecologically realistic contexts, even if they appear so in isolation. The shikimate pathway in RR plants appears to be functionally modified in ways that subtly but consistently redirect metabolic responses under multi-species stress, not causing metabolic breakdown, but altering the balance between generalized and specialized defenses in ways that propagate upward through tritrophic interactions.

### Herbivore performance reflects shifts in defense and nutritional metabolism

Our findings emphasize that BPMV infection and rhizobacteria inoculation have a stronger effect on beetle performance than genotype alone. To address this directly, a full set of differential abundance contrasts was computed (Dataset S2), capturing treatment-induced metabolomic changes relative to the uninfected baseline for both NonRR and RR plants. To aid interpretation, Figs. S1 and S2 summarize the number of DAMs per compound class across treatments for NonRR and RR plants respectively, directly linking treatment-induced metabolic breadth to the herbivore outcomes discussed below.

Despite minimal disruption to global metabolic organization under control conditions, RR plants without any biotic stressors supported higher larval survival than their nonRR counterparts, though larval weight was slightly reduced. This suggests that the RR modification subtly alters plant biochemistry, possibly shifting nutrient ratios or stress-response signaling in a way that favors survival while limiting growth. However, beetle weight increased significantly on BPMV*-*infected plants in both genotypes, indicating a general nutritional enhancement or reduction in deterrent defenses triggered by the viral infection. This was especially evident in NonRR control uninfected plants, which showed coordinated increases in alkaloids, saccharides, and fatty acids, particularly within ME1 and ME10 modules—both associated with amino acid metabolism and carbohydrate abundance. Flavonoid accumulation was less consistent, and although some compounds increased (e.g., isorhamnetin 3-galactoside), known deterrents like kaempferol-3-O-glucoside decreased. These metabolite shifts align with an improved feeding environment for *E. varivestis* as previously reported (Pulido *et al*., 2024). In RR plants, viral infection also triggered notable increases in carbohydrates and isoflavonoids (e.g., biochanin), yet these shifts were accompanied by substantial reductions in flavonoids and shikimate-related compounds. While this still resulted in increased larval weight, the lack of change in survival suggests that the modified EPSPS gene may buffer some nutritional or defensive shifts induced by the virus.

In plants singly inoculated with *Bradyrhizobium japonicum* (Bj), herbivore performance increased across both soybean genotypes, regardless of infection status. Larvae gained more weight and showed higher survival odds, a pattern mirrored by substantial enrichment in carbohydrates (e.g., melibiose, cellobiose, methyl galactoside), amino acids and peptides (e.g., glycine, phenylalanine), and fatty acids (e.g., stearic acid, 12,16-octadecadiynoic acid), which are critical for insect nutrition and growth. These enhancements occurred despite the presence or upregulation of flavonoid and alkaloid defenses such as chrysin, indol-acrylate, and thiadiazole derivatives. While flavonoids are classically associated with herbivore deterrence, their effectiveness against *E. varivestis* may be limited because this species is a legume-specialist that might have physiological tolerance to common soybean flavonoids (Gedling *et al*., 2018). That elevated defensive compounds did not suppress herbivore performance likely reflects the specialist nature of *E. varivestis*: coumestrol, strongly elevated under rhizobacterial inoculation and infection, deters generalist herbivores (Sutherland *et al*., 1980) but has no deterrent effect on this species in bioassays (Burden & Norris, 1992). The beetle’s regurgitant contains detoxification enzymes including cytochrome P450s and glutathione-S-transferases (Gedling *et al*., 2018) that likely neutralize such isoflavonoids, leaving nutritional quality as the dominant driver of performance.

In contrast, the increased abundance of sugars observed in Bj-inoculated plants may act as feeding stimulants for this beetle; monosaccharides and sugar alcohols have been shown to promote feeding in *E. varivestis* (Augustine *et al*., 1964), consistent with our observed performance patterns. In this context, increased larval performance on Bj-inoculated plants reflects both enhanced nutritional quality and a reduced defensive state—an interaction previously noted in non-GM soybean lines (Pulido *et al*., 2024). However, the benefit of *B. japonicum* inoculation was somewhat diminished in RR plants, especially for larval survival. This suggests that the RR genetic background alters microbial signal integration, possibly by reactivating select defense pathways or shifting nutrient allocation in ways that restrict herbivore exploitation.

In contrast to *B. japonicum* inoculation, plants treated with *D. acidovorans* exhibited a markedly different metabolic response. In both soybean varieties, *D. acidovorans* led to significant decrease of several primary and secondary metabolites relative to the uninfected baseline, including key compounds in the amino acid, alkaloid, terpenoid, fatty acid and carbohydrate pathways, signaling a reduction in accessible nutrients for the herbivore, thus explaining the reduced larval weight gain.

The enhanced performance of *E. varivestis* larvae on dual-inoculated (BjDa) plants—particularly under BPMV co-infection— correlates with a distinct metabolic signature characterized by fewer compounds having significantly altered accumulation compared to single-inoculation treatments. While single *B. japonicum* inoculation elicited broader metabolic perturbations, dual inoculation with *B. japonicum* and *D. acidovorans* produced a more selective profile: reduced defensive flavonoids (e.g., kaempferol-3-*O*-glucoside) and alkaloids coincided with elevated carbohydrates (e.g., D-(+)-cellobiose, myo-inositol) and fatty acids (e.g., galactaric acid, palmitic acid). This suggests that *E. varivestis* benefits from subtle, treatment-specific increases in accessible nutrients. The marked larval weight gain on BjDa inoculated plants likely reflects this nutritional facilitation, as fatty acids and carbohydrates directly support insect growth and energy metabolism. These metabolic responses were consistent across genotypes, underscoring that treatment effects—rather than constitutive genotypic differences—drive herbivore outcomes. Our results show that microbial synergism in dual-inoculated (BjDa) treatments may suppress defensive compound synthesis while selectively enriching metabolites that enhance host nutritional quality, creating an ecological niche more favorable to *E. varivestis*. These treatment-specific shifts in plant chemistry not only shaped herbivore performance but may also have implications for virus transmission. For instance, in BPMV-infected RR plants, especially those co-inoculated with rhizobacteria, larvae fed more and survived longer, which could increase the likelihood of virus acquisition and subsequent transmission. Although direct assays were not performed, the combination of enhanced feeding and prolonged vector presence on these plants provides a plausible route for BPMV spread, linking plant metabolic responses to epidemiological outcomes.

As noted in the Methods, plants inoculated with B. japonicum received a nitrogen-free nutrient solution while uninoculated controls were supplemented with inorganic nitrogen, a design that unavoidably introduces nutritional differences between treatments. This constraint is inherent to rhizobial research: providing nitrogen to inoculated plants would suppress nodulation, undermining the purpose of the experiment, and withholding it from controls would impose nutrient stress unrelated to the treatments of interest. The approach mirrors standard practice in comparable studies (Kontopoulou *et al*., 2015; Allito *et al*., 2021; Win *et al*., 2023; Ramula *et al*., 2023). Nonetheless, several lines of evidence suggest that the patterns we report reflect rhizobacterial colonization rather than nitrogen availability alone. First, *D. acidovorans* treatments, which received full nitrogen supplementation, produced metabolic and herbivore performance outcomes that diverged substantially from both uninoculated controls and *B. japonicum*-inoculated plants, indicating that treatment identity rather than nitrogen status drove the observed differences. Second, the genotype-specific effects on larval survival and nodule investment under combined stress were consistent across rhizobacterial treatments in ways that cannot be attributed to a simple nitrogen-enrichment effect. Third, the selective accumulation of specific isoflavonoids and the suppression of broad-spectrum defense modules in RR plants under multi-species stress reflect patterns of metabolic reorientation characteristic of rhizobial symbiosis and biotic signaling, not of nitrogen fertilization. While nitrogen availability remains a factor that cannot be entirely excluded, the overall interpretation is robust: the metabolic and ecological outcomes we document are shaped primarily by the identity and combination of microbial colonizers and their interactions with plant genotype.

### Interactions with rhizobacteria shape plant growth and nodulation under viral stress

BPMV infection consistently reduced shoot and root biomass across genotypes. While co-inoculation with both rhizobacteria mitigated some viral impacts, enhancing plant biomass and reducing symptom severity, nodulation remained compromised. This reduction in nodulation, consistent with previous studies (Orellana, *et al*., 1987), suggests rhizobacteria may optimize nitrogen allocation under viral stress, though the specific mechanisms remain unclear. Notably, RR plants showed reduced nodulation under dual inoculation and viral infection, indicating a trade-off between the RR trait and symbiotic resilience under combined stresses. While single *B. japonicum* inoculation improved root biomass in RR plants, this benefit disappeared under viral infection, implying that the RR background’s subtle influence on symbiosis is context-dependent—mirroring its marginal effects on herbivore performance. These belowground responses align with aboveground herbivore outcomes: dual inoculation increased shoot biomass under BPMV infection, and these plants supported elevated larval weight gain, likely via improved nutritional quality. These results demonstrate that rhizobacteria-virus interactions dominate over GM traits in shaping both plant performance and insect interactions in this system.

## Conclusion

Genotype-dependent differences between RR and NonRR soybean are undetectable under baseline conditions but emerge consistently when plants face concurrent colonization by rhizobacteria, infection by a viral pathogen, and feeding by a specialist herbivore. Under multi-species biotic stress, RR plants shift metabolic investment toward selective isoflavonoid production and lipid remodeling, at the cost of broader defense compound classes, a pattern with direct consequences for herbivore performance and nodulation under combined stress. The primary drivers of ecological outcomes across genotypes were the microbial and viral interactions: rhizobacterial colonization enhanced herbivore performance and partially buffered virus-induced biomass losses, while BPMV consistently increased larval weight gain. The RR transgene modulated the coordination and magnitude of these responses, particularly attenuating the survival benefits that *B. japonicum* inoculation otherwise confers on herbivores.

These findings have a practical implication for how we study and evaluate GM crops. Single-species, unstressed experimental designs, which dominate both research and regulatory contexts, are structurally incapable of detecting effects that only emerge from multi-species interactions. Our results suggest that GM modifications to core metabolic enzymes like EPSPS may have ecologically meaningful consequences that are invisible in simplified assays yet apparent when plants are embedded in communities resembling those of real agricultural fields. Future work should test whether these effects extend to field conditions with naturally diverse microbial communities, and whether GM modifications to other core metabolic enzymes produce similarly context-dependent ecological outcomes.

## Acknowledgements

The authors thank Dr. Kerry Mauck for providing initial feedback on the manuscript. Thomas Dorsey at the New Jersey Department of Agriculture’s Phillip Alampi Beneficial Insect Laboratory kindly provided us with beetles as needed. Seeds were provided by Monsanto Company. H.P. is a recipient of the Fulbright/ COLCIENCIAS fellowship.

## Author contributions

H.P., M.C.M. and C.D.M. conceived and designed the study; H.P. conducted the experiments, acquired and curated data, performed statistical analyses, and drafted the initial manuscript. All authors contributed to revisions and gave final approval for publication.

## Data availability

The complete results of the linear models (limma) are provided as Excel files in the supplementary information. Additional supplementary tables and figures are available in the ETHZ repository https://doi.org/10.3929/ethz-c-000797435. Scripts used for statistical analyses are publicly available at: https://gitlab.ethz.ch/hannierp/soybean_monsanto.git

## Supporting information

| Filename | Brief description |
| --- | --- |
| Supplementary_Information_Pulido_et_al_2026.pdf | Supplementary Tables S1–S14 and Supplementary Figures S1–S2. |
| Dataset_S1_limma_RR_vs_NonRR_metabolomics.xlsx | Limma differential abundance results comparing RR vs. NonRR soybean within each rhizobacteria × virus treatment; includes metabolite IDs, compound names, log2FC, p-values, WGCNA module assignments, and NPClassifier pathway annotations. |
| Dataset_S2_limma_treatment_vs_baseline_metabolomics.xlsx | Limma differential abundance results comparing each treatment against the within-genotype control-uninfected baseline, separately for NonRR and RR plants; same column structure as Dataset S1. |
| Dataset_S3_mzmine_parameters.csv | MZmine3 processing parameters used for LC–MS/MS feature detection, alignment, and isotope filtering; required to reproduce the metabolite feature table from raw data. |
| Dataset_S4_Compound_IDs.xlsx | Compound identification table linking metabolite feature IDs across all extraction phases (LC–MS/MS positive and negative modes, GC–MS) to putative compound names, KEGG identifiers, and NPClassifier annotations used throughout the analyses. |
| Table S1 | Full factorial design combining rhizobacterial inoculation treatments and BPMV infection status applied to both RR and NonRR soybean plants. |
| Table S2 | Summary statistics for the GLMM of larval dry weight; Gaussian model fitted with glmmTMB. |
| Table S3 | Pairwise contrasts for larval dry weight; RR vs. NonRR comparisons and within-genotype treatment effects relative to the control-uninfected baseline, with 95% CIs. |
| Table S4 | Summary statistics for the GLMM of larval survival; binomial model fitted with glmmTMB. |
| Table S5 | Pairwise contrasts for larval survival; odds ratios with 95% CIs for RR vs. NonRR comparisons and within-genotype baseline contrasts. |
| Table S6 | Summary statistics for the GLMM of shoot dry biomass; Gaussian model fitted with glmmTMB. |
| Table S7 | Pairwise contrasts for shoot dry biomass; RR vs. NonRR comparisons and within-genotype baseline contrasts with 95% CIs. |
| Table S8 | Summary statistics for the GLMM of root dry biomass; Gaussian model fitted with glmmTMB. |
| Table S9 | Pairwise contrasts for root dry biomass; RR vs. NonRR comparisons and within-genotype baseline contrasts with 95% CIs. |
| Table S10 | Summary statistics for the GLMM of nodule dry biomass; Gaussian model fitted with glmmTMB. |
| Table S11 | Pairwise contrasts for nodule dry biomass; RR vs. NonRR comparisons and within-genotype baseline contrasts with 95% CIs. |
| Table S12 | Summary statistics for the ordinal GLMM of virus symptom severity; cumulative link model fitted with the ordinal package. |
| Table S13 | Summary statistics for the GLMM of shoot biomass as a function of nodule biomass; Gaussian model fitted with glmmTMB including interactions with rhizobacterial treatment, plant variety, and virus infection. |
| Table S14 | Estimated slopes and 95% CIs for the shoot biomass–nodule biomass relationship within each rhizobacteria × variety × virus subgroup. |
| Figure S1 | Differentially accumulated metabolites (DAMs) per biosynthetic pathway class across rhizobacterial and viral treatments in NonRR plants, relative to the NonRR control-uninfected baseline. |
| Figure S2 | Differentially accumulated metabolites (DAMs) per biosynthetic pathway class across rhizobacterial and viral treatments in RR plants, relative to the RR control-uninfected baseline. |

