## Supplementary material for "Ecological context unmasks cryptic effects of glyphosate tolerance on soybean metabolism and performance of the virus vector *Epilachna varivestis*": Supplementary_Information_Pulido_et_al_2026.pdf

Hannier Pulido[1], Consuelo M. De Moraes[1], Mark C. Mescher[1*]

[1] Department of Environmental Systems Science, ETH Zurich, Zurich 8092, Switzerland

**Table S2. Larval weight gain: summary statistics for GLMM** **2**

**Table S3. Larval weight gain: pairwise contrasts** **2**

**Table S4. Larval survival: summary statistics for GLMM** **3**

**Table S5. Larval survival: pairwise contrasts** **4**

**Table S6. Shoot biomass: summary statistics for GLMM** **4**

**Table S7. Shoot biomass: pairwise contrasts** **5**

**Table S8. Root biomass: summary statistics for GLMM** **6**

**Table S9. Root biomass: pairwise contrasts** **6**

**Table S10. Nodule biomass: summary statistics for GLMM** **7**

**Table S11. Nodule biomass: pairwise contrasts** **7**

**Table S12. Virus symptoms: summary statistics for GLMM** **8**

**Table S13. Shoot-Nodulation regression model: summary statistics for GLMM** **8**

**Table S14. Shoot-Nodulation: Slopes for interaction terms** **9**

**Figure S1. Differential metabolite abundance by pathway in NonRR plants** **10**

**Figure S2. Differential metabolite abundance by pathway in RR plants** **11**

#### Table S1. Factorial design combining rhizobacterial treatments and BPMV infection status

Identical treatment combinations were applied to both Roundup Ready (RR) and conventional (NonRR) soybean plants. Bj: *Bradyrhizobium japonicum*; Da: *Delftia acidovorans*; BjDa: Dual inoculation

| | Rhizobacteria Treatments | | | |
|---|---|---|---|---|
| | Control | B. japonicum | D. acidovorans | BjDa |
| **BPMV-infected** | Control infected | Bj infected | Da infected | BjDa infected |
| **BPMV-uninfected** | Control uninfected | Bj uninfected | Da uninfected | BjDa uninfected |

#### Table S2. Larval weight gain: summary statistics for GLMM

The model was fitted using glmmTMB with the formula `dryWeight_per_larvae_mg ~ (rhi + virus + variety)^2`, employing a Gaussian distribution. Significant effects are highlighted in bold.

| term | estimate | **95% CI** | p-value |
|---|---|---|---|
| rhi | | | |
| Control | — | — | |
| Bj | **2.1** | 0.11, 4.1 | **0.038** |
| Da | -2.3 | -4.9, 0.29 | 0.081 |
| BjDa | **2.5** | 0.53, 4.5 | **0.013** |
| virus | | | |
| uninfected | — | — | |
| infected | **2.4** | 0.17, 4.6 | **0.035** |
| variety | | | |
| NonRR | — | — | |
| RR | -2.0 | -4.0, 0.07 | 0.059 |
| rhi * virus | | | |
| Bj * infected | -0.91 | -3.3, 1.5 | 0.5 |
| Da * infected | 1.4 | -1.5, 4.2 | 0.3 |
| BjDa * infected | -0.43 | -2.8, 1.9 | 0.7 |
| rhi * variety | | | |
| Bj * RR | 1.7 | -0.57, 4.0 | 0.14 |
| Da * RR | 2.0 | -0.81, 4.8 | 0.2 |
| BjDa * RR | 2.2 | -0.05, 4.5 | 0.056 |
| virus * variety | | | |
| infected * RR | -0.22 | -1.6, 1.2 | 0.8 |

#### Table S3. Larval weight gain: pairwise contrasts

(A) Comparisons between NonRR vs. RR for each rhizobacteria and virus treatment combination.
(B) Treatment effects relative to control-uninfected baseline within each genotype. Estimates with 95% CIs are reported for all contrasts. Significant effects are highlighted in bold.

| contrast | estimate | lower.CL | upper.CL | p.value |
|---|---|---|---|---|
| RR Control uninfected - NonRR Control uninfected | -1.9501942 | -4.0018623 | 0.1014738 | 0.062 |

*(continued)*

| contrast | estimate | lower.CL | upper.CL | p.value |
|---|---|---|---|---|
| RR Bj uninfected - NonRR Bj uninfected | -0.2261434 | -1.5946228 | 1.1423359 | 0.743 |
| RR Da uninfected - NonRR Da uninfected | 0.0279914 | -2.1287426 | 2.1847253 | 0.979 |
| RR BjDa uninfected - NonRR BjDa uninfected | 0.2763016 | -1.0621806 | 1.6147837 | 0.682 |
| RR Control infected - NonRR Control infected | -2.1686670 | -4.3819941 | 0.0446602 | 0.055 |
| RR Bj infected - NonRR Bj infected | -0.4446162 | -1.8416428 | 0.9524105 | 0.528 |
| RR Da infected - NonRR Da infected | -0.1904813 | -2.2392508 | 1.8582882 | 0.854 |
| RR BjDa infected - NonRR BjDa infected | 0.0578288 | -1.3231711 | 1.4388287 | 0.934 |

| | | | | |
|---|---|---|---|---|
| **NonRR Control infected - NonRR Control uninfected** | **2.3833314** | **0.1338674** | **4.6327954** | **0.038** |
| **NonRR Bj uninfected - NonRR Control uninfected** | **2.0930909** | **0.0846068** | **4.1015749** | **0.041** |
| **NonRR Bj infected - NonRR Control uninfected** | **3.5621340** | **1.3585875** | **5.7656805** | **0.002** |
| NonRR Da uninfected - NonRR Control uninfected | -2.3061271 | -4.9373331 | 0.3250789 | 0.085 |
| NonRR Da infected - NonRR Control uninfected | 1.4674267 | -1.0925526 | 4.0274059 | 0.258 |
| **NonRR BjDa uninfected - NonRR Control uninfected** | **2.5285417** | **0.5052916** | **4.5517919** | **0.015** |
| **NonRR BjDa infected - NonRR Control uninfected** | **4.4860809** | **2.3201926** | **6.6519692** | **<0.0001** |
| RR Control infected - RR Control uninfected | 2.1648586 | -0.0329273 | 4.3626446 | 0.053 |
| **RR Bj uninfected - RR Control uninfected** | **3.8171417** | **2.1104898** | **5.5237935** | **<0.0001** |
| **RR Bj infected - RR Control uninfected** | **5.0677121** | **3.3572642** | **6.7781599** | **<0.0001** |
| RR Da uninfected - RR Control uninfected | -0.3279415 | -2.5220360 | 1.8661530 | 0.767 |
| **RR Da infected - RR Control uninfected** | **3.2271396** | **1.2411733** | **5.2131059** | **0.002** |
| **RR BjDa uninfected - RR Control uninfected** | **4.7550375** | **3.1052634** | **6.4048116** | **<0.0001** |
| **RR BjDa infected - RR Control uninfected** | **6.4941040** | **4.7525461** | **8.2356618** | **<0.0001** |

### Table S4. Larval survival: summary statistics for GLMM

The model was fitted using glmmTMB with the formula `cbind(larvae_number, 10 - larvae_number) ~ (rhi + virus + variety)^3` on binomial proportion data, employing a logistic regression model. Significant effects are highlighted in bold.

| term | estimate | **95% CI** | p-value |
|---|---|---|---|
| rhi | | | |
| Control | — | — | |
| Bj | **1.5** | 0.44, 2.5 | **0.005** |
| Da | 0.21 | -1.3, 1.8 | 0.8 |
| BjDa | **1.3** | 0.27, 2.3 | **0.013** |
| virus | | | |
| uninfected | — | — | |
| infected | 1.1 | -0.24, 2.4 | 0.11 |
| variety | | | |
| NonRR | — | — | |
| RR | **1.2** | 0.17, 2.3 | **0.023** |
| rhi * virus | | | |
| Bj * infected | -1.3 | -2.8, 0.16 | 0.082 |
| Da * infected | -0.55 | -2.6, 1.5 | 0.6 |
| BjDa * infected | -1.0 | -2.5, 0.47 | 0.2 |
| rhi * variety | | | |
| Bj * RR | **-1.6** | -2.8, -0.41 | **0.009** |
| Da * RR | -1.5 | -3.3, 0.35 | 0.11 |
| BjDa * RR | -1.1 | -2.4, 0.06 | 0.062 |
| virus * variety | | | |

*(continued)*

| term | estimate | **95% CI** | p-value |
|---|---|---|---|
| infected * RR | -1.2 | -2.9, 0.50 | 0.2 |
| rhi * virus * variety | | | |
| Bj * infected * RR | 1.7 | -0.21, 3.6 | 0.081 |
| Da * infected * RR | 1.3 | -1.2, 3.8 | 0.3 |
| BjDa * infected * RR | 1.3 | -0.57, 3.2 | 0.2 |

### Table S5. Larval survival: pairwise contrasts

(A) Comparisons between NonRR vs. RR for each rhizobacteria and virus treatment combination.
(B) Treatment effects relative to control-uninfected baseline within each genotype. Odds ratios with 95% CIs are reported for all contrasts.

| contrast | odds.ratio | lower.CL | upper.CL | p.value |
|---|---|---|---|---|
| RR BjDa infected - NonRR BjDa infected | 1.2058295 | 0.6617151 | 2.1973580 | 0.541 |
| RR BjDa uninfected - NonRR BjDa uninfected | 1.0705697 | 0.5896281 | 1.9438009 | 0.823 |
| RR Da infected - NonRR Da infected | 0.8214360 | 0.2856215 | 2.3624165 | 0.715 |
| RR Da uninfected - NonRR Da uninfected | 0.7727282 | 0.1733417 | 3.4446929 | 0.735 |
| RR Bj infected - NonRR Bj infected | 1.0769263 | 0.5905730 | 1.9638053 | 0.809 |
| RR Bj uninfected - NonRR Bj uninfected | 0.6628268 | 0.3605255 | 1.2186084 | 0.186 |
| RR Control infected - NonRR Control infected | 0.9999963 | 0.2585905 | 3.8670890 | 1 |
| **RR Control uninfected - NonRR Control uninfected** | **3.3703615** | **1.1825523** | **9.6057798** | **0.023** |
| NonRR Control infected - NonRR Control uninfected | 2.9999707 | 0.7863860 | 11.4445385 | 0.108 |
| **NonRR Bj uninfected - NonRR Control uninfected** | **4.2903117** | **1.5465378** | **11.9019235** | **0.005** |
| **NonRR Bj infected - NonRR Control uninfected** | **3.4999808** | **1.2439507** | **9.8475492** | **0.018** |
| NonRR Da uninfected - NonRR Control uninfected | 1.2352785 | 0.2637143 | 5.7862352 | 0.789 |
| NonRR Da infected - NonRR Control uninfected | 2.1304177 | 0.6028070 | 7.5292419 | 0.24 |
| **NonRR BjDa uninfected - NonRR Control uninfected** | **3.6779353** | **1.3091255** | **10.3330107** | **0.013** |
| **NonRR BjDa infected - NonRR Control uninfected** | **4.0526115** | **1.4461457** | **11.3568500** | **0.008** |
| RR Control infected - RR Control uninfected | 0.8901002 | 0.3069435 | 2.5811865 | 0.83 |
| RR Bj uninfected - RR Control uninfected | 0.8437474 | 0.4390717 | 1.6213974 | 0.61 |
| RR Bj infected - RR Control uninfected | 1.1183434 | 0.5999910 | 2.0845178 | 0.725 |
| **RR Da uninfected - RR Control uninfected** | **0.2832143** | **0.1070561** | **0.7492368** | **0.011** |
| RR Da infected - RR Control uninfected | 0.5192326 | 0.2364089 | 1.1404078 | 0.103 |
| RR BjDa uninfected - RR Control uninfected | 1.1682682 | 0.6278477 | 2.1738563 | 0.624 |
| RR BjDa infected - RR Control uninfected | 1.4499212 | 0.7732659 | 2.7186915 | 0.247 |

### Table S6. Shoot biomass: summary statistics for GLMM

The model was fitted using glmmTMB with the formula `shoot_dry_weight ~ (rhi + virus + variety)^3`, employing a Gaussian distribution. Significant effects are highlighted in bold.

| term | estimate | **95% CI** | p-value |
|---|---|---|---|
| rhi | | | |
| Control | — | — | |
| Bj | 0.20 | -0.58, 0.98 | 0.6 |
| Da | **-1.1** | -1.9, -0.32 | **0.006** |
| BjDa | **0.79** | 0.00, 1.6 | **0.049** |

*(continued)*

| term | estimate | **95% CI** | p-value |
|---|---|---|---|
| virus | | | |
| uninfected | — | — | |
| infected | **-1.7** | -2.5, -0.89 | **<0.001** |
| variety | | | |
| NonRR | — | — | |
| RR | 0.36 | -0.42, 1.1 | 0.4 |
| rhi * virus | | | |
| Bj * infected | 0.96 | -0.15, 2.1 | 0.089 |
| Da * infected | 1.0 | -0.09, 2.1 | 0.073 |
| BjDa * infected | -0.34 | -1.5, 0.76 | 0.5 |
| rhi * variety | | | |
| Bj * RR | 0.64 | -0.47, 1.8 | 0.3 |
| Da * RR | 0.28 | -0.83, 1.4 | 0.6 |
| BjDa * RR | -0.60 | -1.7, 0.51 | 0.3 |
| virus * variety | | | |
| infected * RR | -0.30 | -1.4, 0.80 | 0.6 |
| rhi * virus * variety | | | |
| Bj * infected * RR | -1.0 | -2.6, 0.53 | 0.2 |
| Da * infected * RR | 0.05 | -1.5, 1.6 | >0.9 |
| BjDa * infected * RR | -0.05 | -1.6, 1.5 | >0.9 |

### Table S7. Shoot biomass: pairwise contrasts

(A) Comparisons between NonRR vs. RR for each rhizobacteria and virus treatment combination.
(B) Treatment effects relative to control-uninfected baseline within each genotype. Estimates with 95% CIs are reported for all contrasts. Significant effects are highlighted in bold.

| contrast | estimate | lower.CL | upper.CL | p.value |
|---|---|---|---|---|
| RR BjDa infected - NonRR BjDa infected | -0.5915796 | -1.3907002 | 0.2075410 | 0.144 |
| RR BjDa uninfected - NonRR BjDa uninfected | -0.2386610 | -1.0377816 | 0.5604596 | 0.553 |
| RR Da infected - NonRR Da infected | 0.3789000 | -0.4202206 | 1.1780205 | 0.347 |
| RR Da uninfected - NonRR Da uninfected | 0.6387399 | -0.1603807 | 1.4378605 | 0.115 |
| RR Bj infected - NonRR Bj infected | -0.3370403 | -1.1361608 | 0.4620803 | 0.403 |
| **RR Bj uninfected - NonRR Bj uninfected** | **1.0011002** | **0.2019796** | **1.8002208** | **0.015** |
| RR Control infected - NonRR Control infected | 0.0543796 | -0.7447410 | 0.8535002 | 0.892 |
| RR Control uninfected - NonRR Control uninfected | 0.3593000 | -0.4398206 | 1.1584206 | 0.372 |
| **NonRR Control infected - NonRR Control uninfected** | **-1.6700595** | **-2.4691800** | **-0.8709389** | **<0.0001** |
| NonRR Bj uninfected - NonRR Control uninfected | 0.1999400 | -0.5991806 | 0.9990606 | 0.619 |
| NonRR Bj infected - NonRR Control uninfected | -0.5090998 | -1.3082204 | 0.2900207 | 0.208 |
| **NonRR Da uninfected - NonRR Control uninfected** | **-1.1002798** | **-1.8994004** | **-0.3011593** | **0.008** |
| **NonRR Da infected - NonRR Control uninfected** | **-1.7548400** | **-2.5539606** | **-0.9557194** | **<0.0001** |
| NonRR BjDa uninfected - NonRR Control uninfected | 0.7863008 | -0.0128198 | 1.5854213 | 0.054 |
| **NonRR BjDa infected - NonRR Control uninfected** | **-1.2278202** | **-2.0269408** | **-0.4286997** | **0.003** |
| **RR Control infected - RR Control uninfected** | **-1.9749799** | **-2.7741004** | **-1.1758593** | **<0.0001** |
| **RR Bj uninfected - RR Control uninfected** | **0.8417402** | **0.0426196** | **1.6408608** | **0.039** |
| **RR Bj infected - RR Control uninfected** | **-1.2054401** | **-2.0045607** | **-0.4063195** | **0.004** |
| **RR Da uninfected - RR Control uninfected** | **-0.8208400** | **-1.6199605** | **-0.0217194** | **0.044** |
| **RR Da infected - RR Control uninfected** | **-1.7352400** | **-2.5343606** | **-0.9361194** | **<0.0001** |
| RR BjDa uninfected - RR Control uninfected | 0.1883398 | -0.6107808 | 0.9874603 | 0.639 |

*(continued)*

| contrast | estimate | lower.CL | upper.CL | p.value |
|---|---|---|---|---|
| **RR BjDa infected - RR Control uninfected** | **-2.1786998** | **-2.9778204** | **-1.3795793** | **<0.0001** |

### Table S8. Root biomass: summary statistics for GLMM

The model was fitted using glmmTMB with the formula `root_dry_weight ~ (rhi + virus + variety)^3`, employing a Gaussian distribution. Significant effects are highlighted in bold.

| term | estimate | **95% CI** | p-value |
|---|---|---|---|
| rhi | | | |
| Control | — | — | |
| Bj | **-0.55** | -0.90, -0.20 | **0.002** |
| Da | -0.11 | -0.46, 0.24 | 0.6 |
| BjDa | -0.22 | -0.57, 0.13 | 0.2 |
| virus | | | |
| uninfected | — | — | |
| infected | **-0.61** | -0.96, -0.26 | **<0.001** |
| variety | | | |
| NonRR | — | — | |
| RR | 0.02 | -0.33, 0.37 | >0.9 |
| rhi * virus | | | |
| Bj * infected | **0.61** | 0.11, 1.1 | **0.016** |
| Da * infected | -0.06 | -0.55, 0.44 | 0.8 |
| BjDa * infected | -0.06 | -0.56, 0.43 | 0.8 |
| rhi * variety | | | |
| Bj * RR | **0.59** | 0.09, 1.1 | **0.020** |
| Da * RR | 0.05 | -0.45, 0.54 | 0.8 |
| BjDa * RR | -0.15 | -0.64, 0.35 | 0.6 |
| virus * variety | | | |
| infected * RR | -0.28 | -0.77, 0.22 | 0.3 |
| rhi * virus * variety | | | |
| Bj * infected * RR | -0.60 | -1.3, 0.10 | 0.10 |
| Da * infected * RR | 0.43 | -0.27, 1.1 | 0.2 |
| BjDa * infected * RR | 0.11 | -0.59, 0.81 | 0.8 |

### Table S9. Root biomass: pairwise contrasts

(A) Comparisons between NonRR vs. RR for each rhizobacteria and virus treatment combination.
(B) Treatment effects relative to control-uninfected baseline within each genotype. Estimates with 95% CIs are reported for all contrasts. Significant effects are highlighted in bold.

| contrast | estimate | lower.CL | upper.CL | p.value |
|---|---|---|---|---|
| RR BjDa infected - NonRR BjDa infected | -0.3009600 | -0.6577592 | 0.0558391 | 0.097 |
| RR BjDa uninfected - NonRR BjDa uninfected | -0.1328608 | -0.4896599 | 0.2239384 | 0.46 |
| RR Da infected - NonRR Da infected | 0.2171206 | -0.1396786 | 0.5739197 | 0.229 |
| RR Da uninfected - NonRR Da uninfected | 0.0642795 | -0.2925197 | 0.4210786 | 0.72 |
| RR Bj infected - NonRR Bj infected | -0.2697604 | -0.6265595 | 0.0870388 | 0.136 |
| **RR Bj uninfected - NonRR Bj uninfected** | **0.6023403** | **0.2455412** | **0.9591395** | **0.001** |

*(continued)*

| contrast | estimate | lower.CL | upper.CL | p.value |
|---|---|---|---|---|
| RR Control infected - NonRR Control infected | -0.2608206 | -0.6176198 | 0.0959786 | 0.149 |
| RR Control uninfected - NonRR Control uninfected | 0.0159204 | -0.3408787 | 0.3727196 | 0.929 |
| **NonRR Control infected - NonRR Control uninfected** | **-0.6104595** | **-0.9672587** | **-0.2536604** | **0.001** |
| **NonRR Bj uninfected - NonRR Control uninfected** | **-0.5461398** | **-0.9029390** | **-0.1893407** | **0.003** |
| **NonRR Bj infected - NonRR Control uninfected** | **-0.5467793** | **-0.9035785** | **-0.1899801** | **0.003** |
| NonRR Da uninfected - NonRR Control uninfected | -0.1051992 | -0.4619984 | 0.2515999 | 0.558 |
| **NonRR Da infected - NonRR Control uninfected** | **-0.7725598** | **-1.1293590** | **-0.4157607** | **<0.0001** |
| NonRR BjDa uninfected - NonRR Control uninfected | -0.2244192 | -0.5812184 | 0.1323799 | 0.213 |
| **NonRR BjDa infected - NonRR Control uninfected** | **-0.8962596** | **-1.2530587** | **-0.5394604** | **<0.0001** |
| **RR Control infected - RR Control uninfected** | **-0.8872005** | **-1.2439997** | **-0.5304014** | **<0.0001** |
| RR Bj uninfected - RR Control uninfected | 0.0402801 | -0.3165191 | 0.3970792 | 0.822 |
| **RR Bj infected - RR Control uninfected** | **-0.8324601** | **-1.1892593** | **-0.4756610** | **<0.0001** |
| RR Da uninfected - RR Control uninfected | -0.0568402 | -0.4136393 | 0.2999590 | 0.751 |
| **RR Da infected - RR Control uninfected** | **-0.5713597** | **-0.9281588** | **-0.2145605** | **0.002** |
| **RR BjDa uninfected - RR Control uninfected** | **-0.3732004** | **-0.7299996** | **-0.0164013** | **0.041** |
| **RR BjDa infected - RR Control uninfected** | **-1.2131400** | **-1.5699392** | **-0.8563409** | **<0.0001** |

### Table S10. Nodule biomass: summary statistics for GLMM

The model was fitted using glmmTMB with the formula `nodules_dry_weight ~ (rhi + virus + variety)^3`, employing a Gaussian distribution. Significant effects are highlighted in bold.

| term | estimate | **95% CI** | p-value |
|---|---|---|---|
| rhi | | | |
| Bj | — | — | |
| BjDa | -0.01 | -0.10, 0.09 | >0.9 |
| virus | | | |
| uninfected | — | — | |
| infected | -0.05 | -0.15, 0.04 | 0.3 |
| variety | | | |
| NonRR | — | — | |
| RR | -0.01 | -0.10, 0.08 | 0.8 |
| rhi * virus | | | |
| BjDa * infected | -0.03 | -0.16, 0.10 | 0.7 |
| rhi * variety | | | |
| BjDa * RR | -0.03 | -0.16, 0.10 | 0.7 |
| virus * variety | | | |
| infected * RR | -0.04 | -0.17, 0.09 | 0.6 |
| rhi * virus * variety | | | |
| BjDa * infected * RR | -0.03 | -0.22, 0.16 | 0.8 |

### Table S11. Nodule biomass: pairwise contrasts

(A) Comparisons between NonRR vs. RR for each rhizobacteria and virus treatment combination.
(B) Treatment effects relative to control-uninfected baseline within each genotype. Estimates with 95% CIs are reported for all contrasts. Significant effects are highlighted in bold.

| contrast | estimate | lower.CL | upper.CL | p.value |
|---|---|---|---|---|
| **RR BjDa infected - NonRR BjDa infected** | **-0.1098197** | **-0.2068220** | **-0.0128174** | **0.028** |
| RR BjDa uninfected - NonRR BjDa uninfected | -0.0400998 | -0.1371021 | 0.0569025 | 0.406 |
| RR Bj infected - NonRR Bj infected | -0.0498003 | -0.1468026 | 0.0472020 | 0.303 |
| RR Bj uninfected - NonRR Bj uninfected | -0.0103799 | -0.1073822 | 0.0866224 | 0.829 |
| NonRR Bj infected - NonRR Bj uninfected | -0.0543597 | -0.1513620 | 0.0426425 | 0.262 |
| NonRR BjDa uninfected - NonRR Bj uninfected | -0.0050603 | -0.1020626 | 0.0919420 | 0.916 |
| NonRR BjDa infected - NonRR Bj uninfected | -0.0863602 | -0.1833625 | 0.0106421 | 0.079 |
| RR Bj infected - RR Bj uninfected | -0.0937801 | -0.1907824 | 0.0032222 | 0.058 |
| RR BjDa uninfected - RR Bj uninfected | -0.0347802 | -0.1317825 | 0.0622221 | 0.47 |
| **RR BjDa infected - RR Bj uninfected** | **-0.1858000** | **-0.2828023** | **-0.0887978** | **0.00047** |

### Table S12. Virus symptoms: summary statistics for GLMM

Symptom severity was modeled using cumulative link regression (ordinal::clm) with formula `virus_symptoms ~ (rhi + variety)` and logit link.

| term | estimate | **95% CI** | p-value |
|---|---|---|---|
| rhi | | | |
| Control | — | — | |
| Bj | **-3.7** | -6.8, -1.5 | **0.003** |
| Da | **-3.6** | -6.7, -1.5 | **0.004** |
| BjDa | -0.91 | -4.1, 1.6 | 0.5 |
| variety | | | |
| NonRR | — | — | |
| RR | -0.23 | -1.6, 1.1 | 0.7 |

### Table S13. Shoot-Nodulation regression model: summary statistics for GLMM

The model was fitted using glmmTMB with the formula `shoot_dry_weight ~ nodules_dry_weight * (rhi + variety + virus)`, employing a Gaussian distribution. Significant effects are highlighted in bold.

| term | estimate | **95% CI** | p-value |
|---|---|---|---|
| nodules_dry_weight | 3.7 | -1.2, 8.7 | 0.13 |
| rhi | | | |
| Bj | — | — | |
| BjDa | -0.08 | -1.0, 0.87 | 0.9 |
| variety | | | |
| NonRR | — | — | |
| RR | 0.41 | -0.57, 1.4 | 0.4 |
| virus | | | |
| uninfected | — | — | |
| infected | **-2.5** | -3.8, -1.2 | **<0.001** |
| nodules_dry_weight * rhi | | | |
| nodules_dry_weight * BjDa | -0.17 | -3.9, 3.6 | >0.9 |
| nodules_dry_weight * variety | | | |
| nodules_dry_weight * RR | -0.18 | -4.0, 3.6 | >0.9 |

*(continued)*

| term | estimate | **95% CI** | p-value |
|---|---|---|---|
| nodules_dry_weight * virus | | | |
| nodules_dry_weight * infected | **5.6** | 0.77, 10 | **0.023** |

#### Table S14. Shoot-Nodulation: Slopes for interaction terms

(A) Comparisons between NonRR vs. RR for each rhizobacteria and virus treatment combination.
(B) Treatment effects relative to control-uninfected baseline within each genotype. Estimates with 95% CIs are reported for all contrasts. Significant effects are highlighted in bold.

| treatment | Slope (beta) | lower.CL | upper.CL | p.value |
|---|---|---|---|---|
| NonRR Bj uninfected | 3.75 | -1.36 | 8.85 | 0.14 |
| NonRR BjDa uninfected | 3.58 | -2.33 | 9.48 | 0.23 |
| RR Bj uninfected | 3.57 | -1.38 | 8.53 | 0.15 |
| RR BjDa uninfected | 3.40 | -1.70 | 8.51 | 0.18 |
| **NonRR Bj infected** | **9.31** | **6.11** | **12.51** | **<0.0001** |
| **NonRR BjDa infected** | **9.14** | **5.15** | **13.14** | **<0.0001** |
| **RR Bj infected** | **9.14** | **5.17** | **13.11** | **<0.0001** |
| **RR BjDa infected** | **8.97** | **5.20** | **12.74** | **<0.0001** |

#### Figure S1. Differential metabolite abundance by pathway in NonRR plants

Differentially accumulated metabolites (DAMs) by biosynthetic pathway across rhizobacterial and viral treatments, relative to the NonRR control-uninfected baseline. Bars represent the count of metabolites significantly altered ($p$ value $< 0.05$) in each pathway class, with colors indicating direction of change (blue: elevated; green: reduced).

NonRR: near-isogenic non-modified soybean; RR: Roundup Ready soybean; Bj: *Bradyrhizobium japonicum*-inoculated; Da: *Delftia acidovorans*-inoculated; BjDa: dual-inoculated.

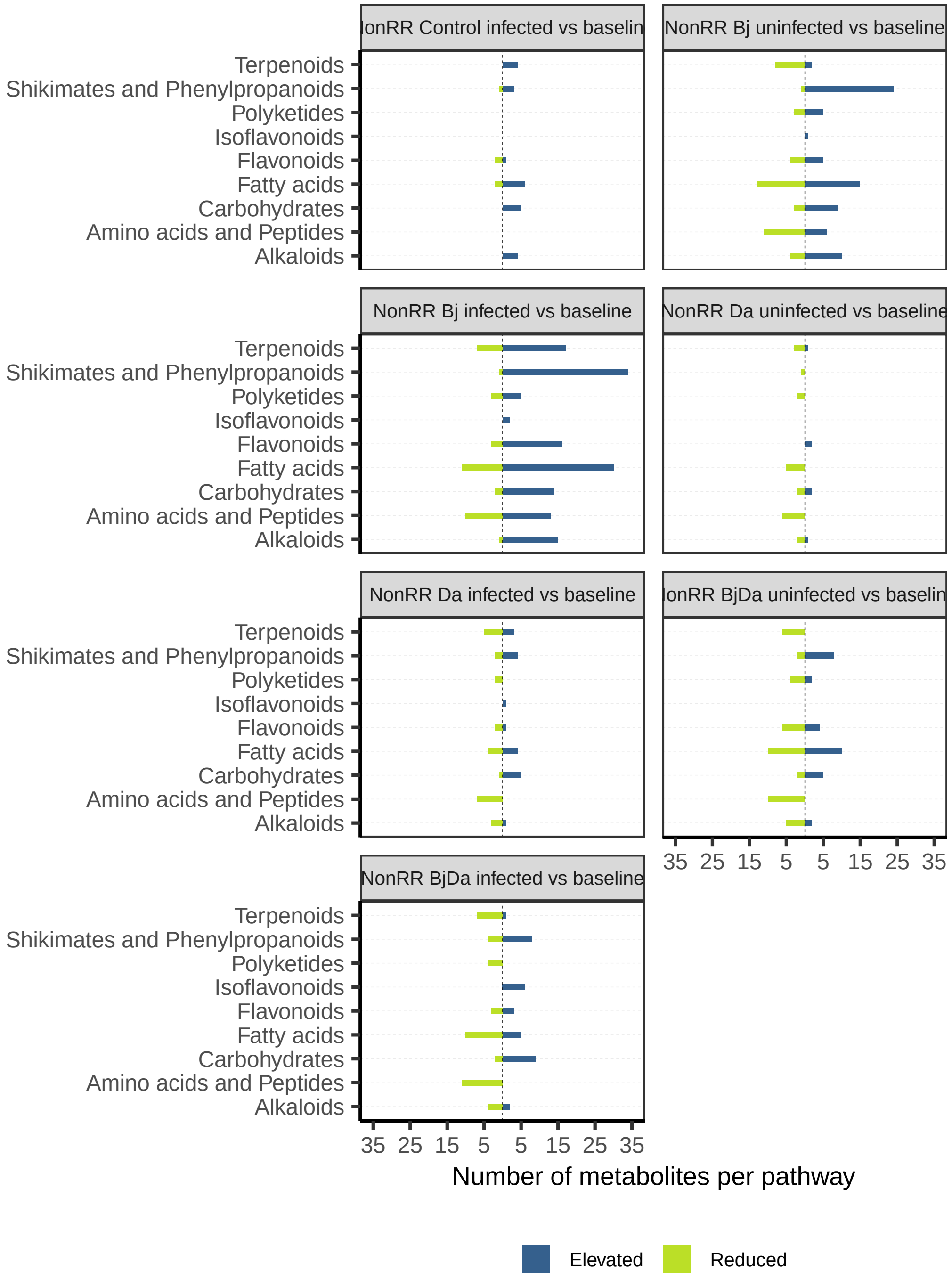

### Figure S2. Differential metabolite abundance by pathway in RR plants

Differentially accumulated metabolites (DAMs) by biosynthetic pathway across rhizobacterial and viral treatments, relative to the RR control-uninfected baseline. Bars represent the count of metabolites significantly altered ($p$ value $< 0.05$) in each pathway class, with colors indicating direction of change (blue: elevated; green: reduced).

NonRR: near-isogenic non-modified soybean; RR: Roundup Ready soybean; Bj: *Bradyrhizobium japonicum*-inoculated; Da: *Delftia acidovorans*-inoculated; BjDa: dual-inoculated.

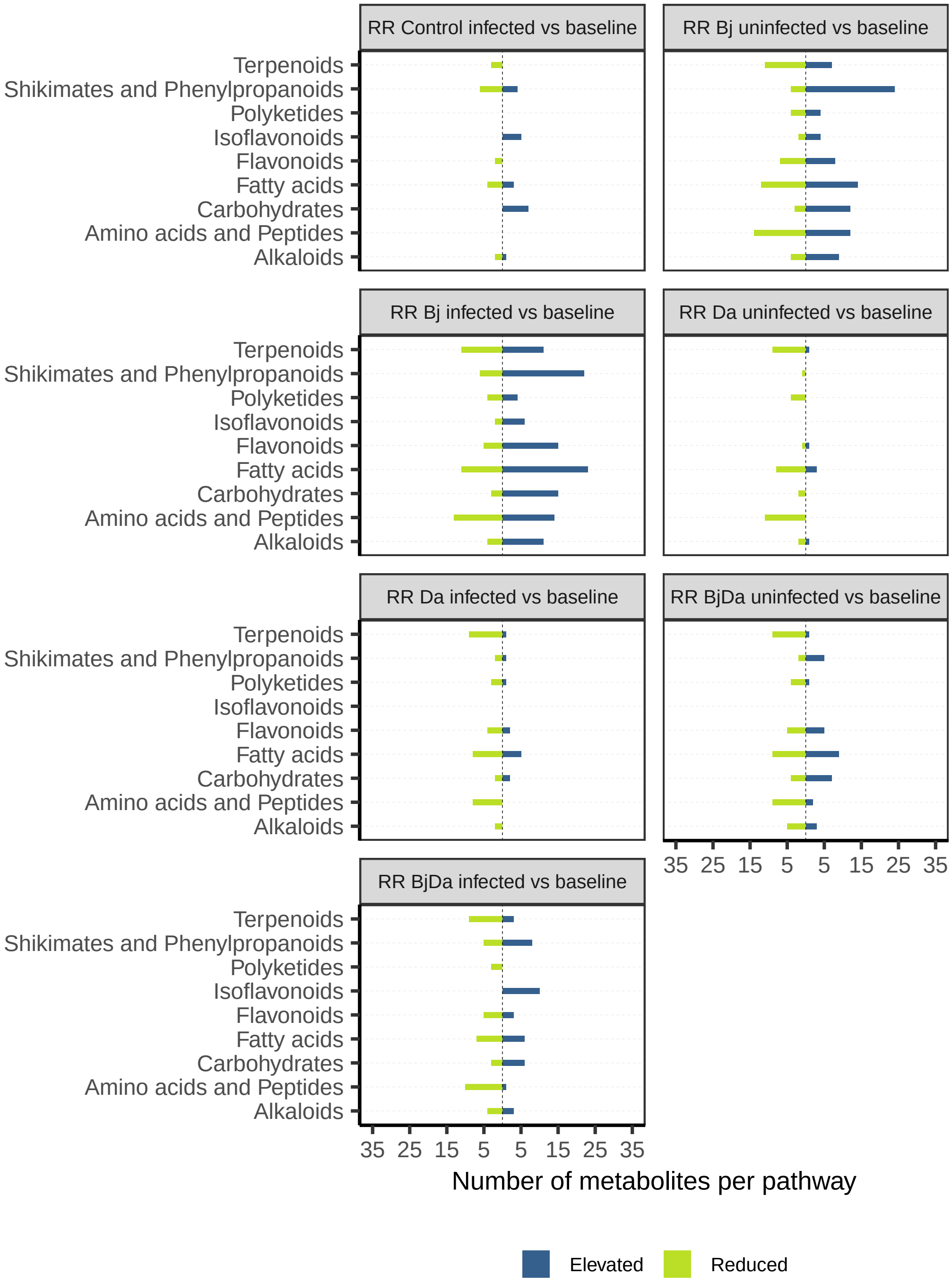
